# Molecular determinants of protein pathogenicity at the single-aggregate level

**DOI:** 10.1101/2024.11.12.623279

**Authors:** Agnieszka Urbanek, Emma F. Garland, Emily E. Prescott, Marianne C. King, Anna Olerinyova, Hollie E. Wareing, Nia Georgieva, Ellie L. Bradshaw, Svetomir B. Tzokov, Alexander Knight, Alexander I. Tartakovskii, Tarja Malm, J Robin Highley, Suman De

**Affiliations:** Sheffield Institute for Translational Neuroscience, Division of Neuroscience, University of Sheffield, Sheffield, S10 2HQ, UK; Cryo-Electron Microscopy Facility, School of Biosciences, University of Sheffield, Sheffield, S10 2TN, UK; Department of Physics and Astronomy, University of Sheffield, Sheffield, S3 7RH, UK; A.I. Virtanen Institute for Molecular Sciences, University of Eastern Finland, Kuopio, 70211, Finland; Neuroscience Institute, University of Sheffield, Sheffield, S10 2TN, UK; Healthy Lifespan Institute, University of Sheffield, Western Bank, Sheffield, S10 2TN, UK

## Abstract

Determining the structure-function relationships of protein aggregates is a fundamental challenge in biology. These aggregates, whether formed *in vitro*, within cells, or in living organisms, present significant heterogeneity in their molecular features such as size, structure, and composition, making it difficult to determine how their structure influences their functions. Interpreting how these molecular features translate into functional roles is crucial for understanding cellular homeostasis and the pathogenesis of various debilitating diseases like Alzheimer’s and Parkinson’s. In this study, we introduce a bottom-up approach to explore how variations in protein aggregates’ size, composition, post-translational modifications and point mutations profoundly influence their biological functions. Applying this method to Alzheimer’s and Parkinson’s associated proteins, we uncover the mechanism of novel disease-relevant pathways and demonstrate how subtle alterations in composition and morphology can shift the balance between healthy and pathological states. Our findings establish a broadly applicable framework for investigating protein dysfunctions in various proteinopathies.

## Introduction

Protein aggregates are basic components of cells, executing essential biochemical processes and orchestrating the smooth functioning of life at the molecular level. The function of these aggregates hinges on the precise and regulated interactions among their molecular components. When the delicate balance of these interactions is disrupted, aberrant protein assemblies can form. These malformations are not merely structural anomalies, they often lead to impaired functionality and contribute to debilitating diseases known as proteinopathies, such as Alzheimer’s (AD) and Parkinson’s (PD)^1,2^ diseases. In these conditions, proteins aggregate into higher-order assemblies in an unregulated manner, causing cellular dysfunctions and eventually cell death. These aberrant assemblies generally differ from their healthy counterparts in one or more key molecular aspects such as size, composition, mutation, and post-translational modification (PTM) that are key drivers in regulating their functions^3,4^. These subtle, yet significant, differences are essential for understanding the disease-related functions of these pathological aggregates and their impact on the progression of proteinopathies.

Structure-function analysis of protein aggregates in native environments like cells, tissues, and biofluids is challenging due to sample complexity. Traditional methods utilizing recombinant proteins to create higher-order structures like oligomers and fibrils provides a more controlled approach for functional measurement. However, protein self-assembly process is highly heterogeneous, even under controlled conditions, producing species with varying sizes, shapes, aggregation states, and compositions that complicate structure-function analysis^5–7^. Proteins like amyloid-beta (Aβ), which is linked to AD^8^, and alpha-synuclein (αSyn), associated with PD^9^, are not toxic in their monomeric forms but become harmful as they aggregate into higher order structures^1,10^. Oligomeric aggregates, forming as intermediates on the pathway to insoluble fibrils, are particularly toxic and contribute to neurodegeneration by damaging neurons and glial cells^1,2,10–12^. These intermediates are transient, exists in much lower concentrations than monomers or fully aggregated fibrils at any stage of aggregation complicating their functional characterization^7,13,14^. Techniques such as ultracentrifugation^15^, sucrose gradient fractionation^7^, and capturing the temporal evolution of the aggregation pathway^7,16^ are employed to enrich specific types of species, but often result in mixed populations and lack consistency in producing samples with well-defined sizes or compositions for reproducible analysis.

Recognizing the limitations, we developed a bottom-up approach to create protein aggregates and complexes that are uniform in size and composition, enabling us to correlate these characteristics with their biological functions at the single-aggregate level. Our approach involves covalently attaching proteins to nanospheres of known size, which then serve as a platform for additional monomeric proteins to self-assemble into aggregates of consistent size and composition. This uniformity allows for systematic investigation into how subtle variations in size, composition, and critical factors like PTM and missense mutations impact the function of protein aggregates. By precisely controlling these attributes at a single-particle level, we dissect the effects of these molecular drivers on the overall disease-related functions of the protein assemblies.

## Results

### Preparation and characterization of size-controlled protein aggregates at the single-particle level

We began our study by determining the size of diffusible Aβ aggregates present in the brain tissue of AD patients. Our goal was to estimate the size heterogeneity of Aβ aggregates present in human brains, and then emulate these sizes of Aβ aggregates in our experiments. These diffusible Aβ aggregates contribute to disease progression by damaging neuronal and glial cells^17,18^. We employed immunohistology with the pan-Aβ 4G8 antibody to identify amyloid plaques in post-mortem tissue from the prefrontal cortex of three AD patients **(Fig. 1A)**, a region typically marked by significant amyloid deposition and neuronal loss. After confirming the presence of plaques, we selected tissues from the same patients that had been flash-frozen instead of formalin-fixed for further analysis. We extracted diffusible Aβ aggregates from these frozen tissues. To measure the size distribution of Aβ species, we employed direct stochastic optical reconstruction microscopy (dSTORM)^19^ using the same 4G8 antibody. This antibody-based super-resolution imaging technique exceeds the capabilities of conventional microscopy, enabling the size measurement of aggregates smaller than diffraction limit of light (200-250nm) and the specific identification of Aβ-containing species within complex tissue extracts **(Fig. 1B)**. The dSTORM imaging results revealed a diverse range of Aβ aggregates, most of them spanning between 35-800nm **(Fig. 1C)**. Due to the resolution limitations of dSTORM imaging in our system, we could not resolve any aggregates smaller than 35nm, setting this as the minimum measurable size.

**Figure 1.**
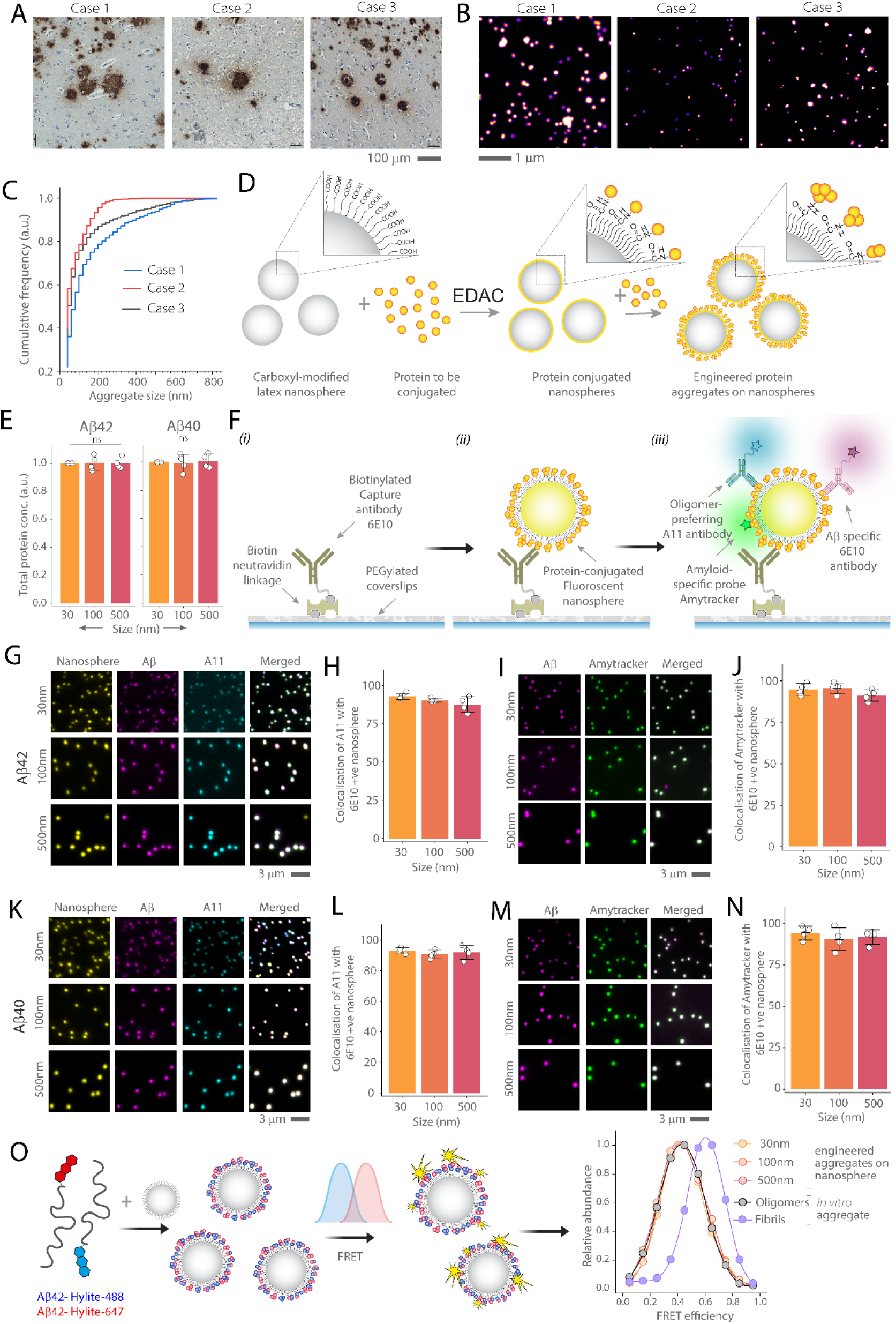
Preparation and characterisation of different sized Aβ40 and Aβ42 aggregates. **(A)** Representative immunohistology images obtained using the Aβ-specific 4G8 antibody shows Aβ plaques in the prefrontal cortex post-mortem tissue from three AD patients. **(B)** Representative dSTORM images show diffusible Aβ aggregates from frozen cortical extracts of the same AD patients detected with the Alexa-647 labelled 4G8 antibody. **(C)** Cumulative size distributions of Aβ aggregates from three patients were measured from dSTORM images. **(D)** Illustration of protein aggregate engineering on the surface of carboxyl-modified latex nanospheres. **(E)** BCA assay measures the protein load on 30nm, 100nm, and 500nm nanospheres, and all data are normalized to the 30nm nanosphere. **(F)** Stepwise protocol for the SiMPull assay for characterizing protein-conjugated nanospheres. **(G-N)** Three-color images of fluorescent nanospheres of various sizes (30nm, 100nm, 500nm) conjugated with Aβ42 **(G, I)** or Aβ40 **(K, M)**. Aggregates are captured using 10nM biotinylated 6E10 antibody and imaged with Alexa 647 labelled 6E10 antibody (1nM) in conjunction with oligomer-specific A11 antibody (1nM) **(G, K)** or the amyloid-specific probe Amytracker (20nM) **(I, M)**. Quantification of three-color colocalization between Aβ42 **(H, J)** or Aβ40 **(L, N)** coupled nanosphere with A11 antibody **(H, L)** or Amytracker **(J, N)**. **(O)** Schematic representation of the FRET assay used to measure the aggregation state of Aβ42 aggregates engineered on nanospheres and compared with in vitro prepared oligomers and fibrils. FRET efficiency data are normalized to the maximum efficiency observed for each condition. Data **(E, H, J, L, N)** are plotted as the mean and standard deviation of four independent replicates. Statistical analyses were performed using one-way ANNOVA with post-hoc Tukey mean comparison. *P < 0.05, **P < 0.01, ***P < 0.001, ns, non-significant (P ≥ 0.05).

Building on these insights, we engineered Aβ40 and Aβ42 aggregates of three distinct sizes: 30nm, 100nm, and 500nm, representing the broad spectrum of Aβ aggregate sizes found in human neural tissue. To achieve this, we first conjugated monomeric Aβ to carboxyl-modified latex nanospheres of 30nm, 100nm, and 500nm **(SI Fig. 1)** and then allowed the protein to aggregate on the surface of each nanosphere. We used EDAC (1-Ethyl-3-(3-dimethylaminopropyl) carbodiimide) as a crosslinker to facilitate covalent attachment of lysine residues of Aβ to the carboxylic acid groups of the nanospheres **(Fig. 1D)**. We calculated the surface area of the nanospheres and added the required amount of Aβ monomer to achieve covalent attachment, considering each monomer diameter is 1nm^20^. To maintain a consistent total surface area across different sizes of nanospheres, we adjusted the number of nanospheres used, while utilizing the same amount of protein for each conjugation. In the next step, we added five times more monomeric protein than the surface area coverage to the protein-coated nanospheres to facilitate the formation of protein aggregates on the surface.

To characterize the engineered protein aggregate nanospheres, we performed Bicinchoninic acid (BCA) **(Fig. 1E)** and Meso Scale Discovery (MSD) **(SI Fig. 2)** assays to verify uniform protein loading across nanospheres of different sizes. Both results confirmed no significant variation across all sizes for both Aβ40 and Aβ42. The MSD assay also showed that more than ∼90% of monomeric proteins are aggregated. To confirm that Aβ aggregates are formed on the surface of the nanospheres the nanospheres, we used the Single-molecule Pull Down (SiMPull) assay^4,21^ **(Fig. 1F),** a method that allows direct visualization and characterization of protein complexes at the single-molecule level. In this assay, we used a Aβ-specific biotinylated 6E10 antibody to capture Aβ aggregates, which were prepared either via conventional *in vitro* aggregation or engineered onto nanosphere surfaces. To confirm that the nanospheres were coated with Aβ aggregates rather than monomers, we introduced a combination of imaging probes: Alexa Fluor 647-labeled 6E10 antibodies along with either Alexa Fluor 561-labeled A11 antibodies, which target oligomeric aggregates^22^, or Amytracker probes^23^ that specifically bind to β-sheet rich aggregates and not to monomers. We used wide-field epi-fluorescence imaging to evaluate the colocalization of the fluorescent nanospheres, 6E10 antibodies and either A11 antibodies or Amytracker probes, across different conditions for both Aβ42 **(Fig. 1G-J)** and Aβ40 **(Fig. 1K-N)**. The mean colocalization across all nanosphere sizes and conditions exceeded 90%, confirming the effectiveness of our conjugation method and verifying that the aggregates formed on the nanospheres as intended. As a control, we used scrambled Aβ42, which shares the same amino acid composition with native Aβ42 but is arranged in a scrambled sequence that does not aggregate. When we conjugated the FAM-labelled scrambled Aβ to the nanospheres, it did not bind to the oligomer-specific A11 antibody **(SI Fig. 3)**. In addition, we conducted a Förster Resonance Energy Transfer (FRET) assay at the single-particle level to study the conformation states **(Fig. 1O)**. For this experiment, we prepared aggregates - both *in vitro* and those engineered on the nanosphere surface - by combining Aβ42 labelled with a FRET donor (HiLyte 488) and an acceptor (HiLyte 647) in a 1:1 molar ratio. This assay was used to determine the relative aggregation state by analyzing FRET efficiencies^24^. Our results revealed that the FRET efficiencies of the engineered aggregates closely matched those of Aβ aggregates formed at the lag phase of *in vitro* aggregation (**Fig. 1O**), which primarily consist of oligomeric species^4,7^ **(SI Fig. 4)**. However, the FRET efficiencies of the engineered aggregates were not aligned with those of Aβ fibrils formed at the plateau phase of aggregation (**Fig. 1O**), confirming that our method effectively mimics the conformation state of oligomeric Aβ and not fibrils. Additionally, we used we used the combination of atomic force microscopy and scattering-type, scanning near-field optical microscopy to visualize the protein coating on the nanospheres **(SI Fig. 5).** While the protein layer was visible, we observed that the nanospheres are clumped due to the solvent-free conditions under which experiments were performed.

### Size-dependent uptake and cytokine secretion by iMGLs in response to Aβ aggregates

After successfully engineering Aβ aggregates of different sizes, we aimed to understand how sizes influence their disease-relevant functions. To investigate this, we focused on the clearance of Aβ aggregates by microglia and the resulting inflammatory responses. Microglia undergo phenotypic activation upon exposure to Aβ aggregates, leading to the secretion of inflammatory cytokine; a key pathological mechanism in AD^25–27^. We used human induced pluripotent stem cell (iPSC)-derived microglia-like cells (iMGLs) for this study. The identities of these iMGLs were previously confirmed by RT-PCR and whole-transcriptome analysis^28^. We exposed these iMGLs to Aβ aggregates for one hour to measure their uptake and assessed the resulting inflammatory activation by measuring the concentrations of three key pro-inflammatory cytokine secreted by iMGLs in the cell culture media: interleukin-1β (IL-1β), interleukin-6 (IL-6), and tumour necrosis factor-alpha (TNF-α) **(Fig. 2A)**. These cytokine are shown to drive inflammation, cause neuronal damage, and ultimately contribute to neurodegeneration in AD^25,29^.

**Figure 2.**
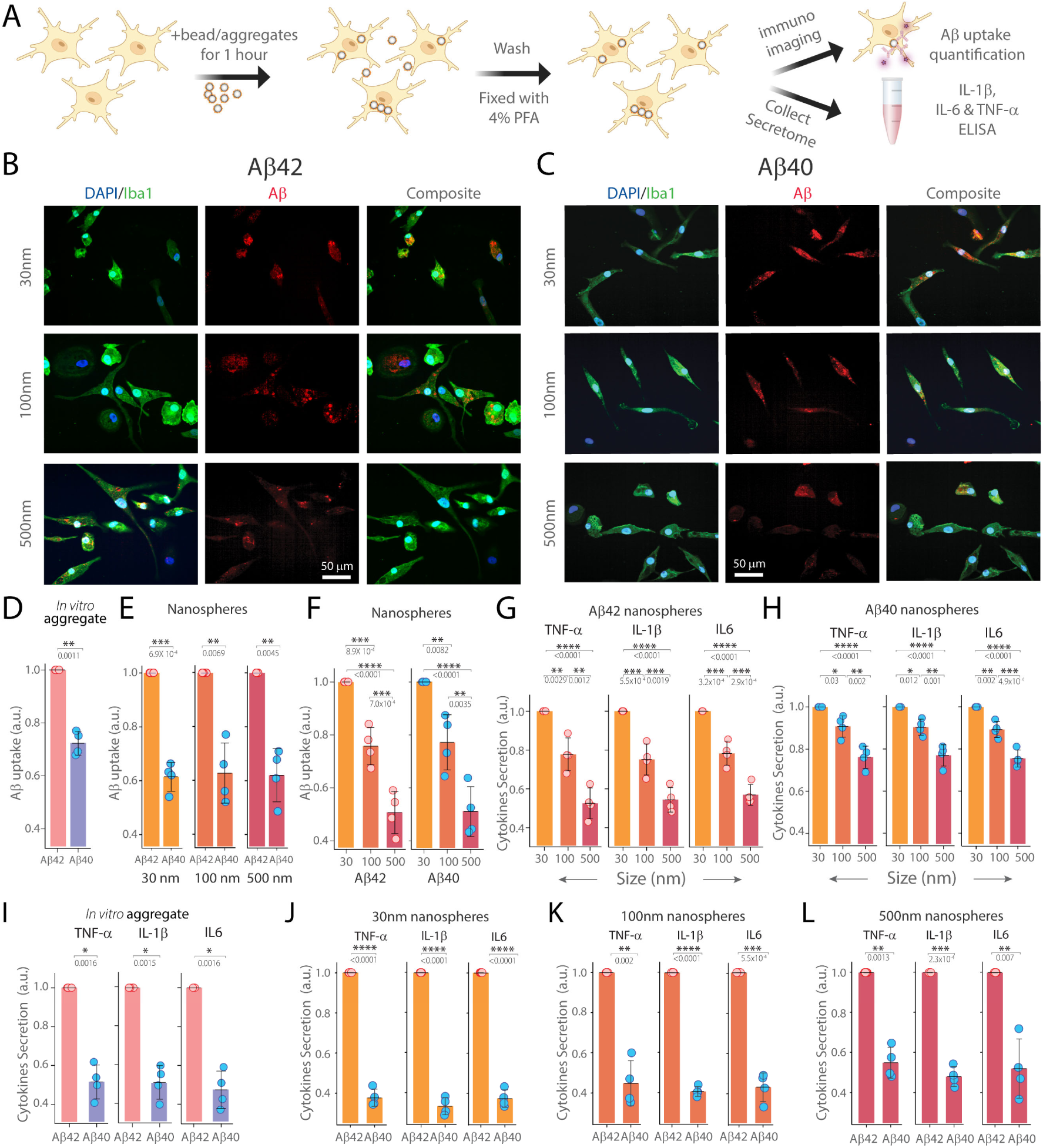
Aβ40 and Aβ42 uptake and subsequent cytokine secretion by iMGLs. **(A)** Schematic of the assessing Aβ uptake by iMGLs and measuring cytokine levels (IL-1β, IL-6, TNF-α) in the media using ELISA. **(B-C)** Representative images of iMGLs (stained with DAPI/Iba1) after uptake of **(B)** Aβ42 and **(C)** Aβ40 aggregates conjugated to nanospheres of 30nm, 100nm, and 500nm. The Aβ internalization was quantified using Aβ-specific 6E10 antibody. **(D-E)** Quantification of Aβ uptake by iMGLs incubated with **(D)** *in vitro* prepared aggregates of Aβ42 and Aβ40 and **(E)** Different sized Aβ conjugated to nanospheres (30nm, 100nm, 500nm). (Units of uptake = integrated fluorescence of sample divided by the integrated fluorescence of corresponding Aβ42 aggregates). **(F)** Comparison of Aβ internalization by iMGLs incubated with Aβ42 and Aβ40 conjugated to nanospheres of different sizes. (Units of uptake = integrated fluorescence of sample divided by the integrated fluorescence of corresponding Aβ aggregates prepared on 30nm nanosphere). **(G-H)** Measurement of cytokine secretion (TNF-α, IL-1β, IL-6) by iMGLs after 1 hour incubation with **(G)** Aβ42 and **(H)** Aβ40 conjugated nanospheres of different sizes (30nm, 100nm, 500nm). Cytokine release was calculated as the cytokine response to the sample divided by the cytokine response to the corresponding Aβ aggregates prepared on 30nm nanospheres **(D, E)** or Aβ aggregates engineered on 30nm nanospheres **(F)** for each replicate. **(I-L)** Comparison of cytokine secretion (TNF-α, IL-1β, IL-6) by iMGLs following incubation with **(I)** *in vitro* prepared Aβ42 and Aβ40 aggregates or **(J)** 30nm, **(K)** 100nm, or **(L)** 500nm Aβ42 and Aβ40 conjugated nanospheres. Units of cytokine release are defined as the total cytokine measured in the media in response to the sample divided by the total cytokine measured in response to corresponding Aβ42 aggregates. Data presented as mean ± standard deviation across four biological replicates. Statistical significance was calculated via unpaired two-sample t-test (D) or one-way ANOVA with Tukey’s post-hoc test (E-L). *P < 0.05, **P < 0.01, ***P < 0.001, ns - non-significant (P ≥ 0.05).

Our results showed that iMGLs were capable of taking up all sizes and isoforms of Aβ aggregates, but to varying extents (**Fig. 2B-C**). We observed that engineered Aβ aggregates on nanospheres were internalized by iMGLs by two orders of magnitude more compared to nanospheres alone (**SI Fig. 6**). Aβ42 aggregates were taken up more by the iMGLs than Aβ40, across both *in vitro* prepared and engineered aggregates on all sizes of nanosphere surfaces (**Fig. 2D-E**). The similarities between *in vitro* prepared and engineered aggregates further validate our methodology. We found that this uptake efficiency of iMGLs is size-dependent, with smaller 30nm aggregates being cleared twice as effectively as larger 500nm aggregates (**Fig. 2F**). We also observed that cytokine release, similar to internalization, is size-dependent for both Aβ42 and Aβ40, with smaller aggregates inducing a stronger response compared to larger ones (**Fig. 2G-H**). Analysis of pro-inflammatory cytokine in the iMGLs media demonstrated that Aβ42 triggered more secretion of TNF-α, IL-1β, and IL-6 than Aβ40, for both types of species - *in vitro* prepared aggregates (**Fig. 2I**) and aggregates engineered on nanosphere surface for all sizes (**Fig. 2J-L**). As a control, we used nanospheres conjugated with scrambled Aβ42 and observed minimal uptake compared to native Aβ42 (**SI Fig. 7**), consistent with previous studies^30^.

### Aggregate size regulates uptake mechanisms and cytokine secretion in iMGLs

Since we found that both uptake and cytokine secretion depend on the size of Aβ aggregates, we investigated which pathways of microglial internalization and subsequent cytokine release are influenced by the size of Aβ. We focused on the effects of TLR-4 (Toll-like receptor 4) inhibition on iMGLs’ responses to differently sized Aβ aggregates. TLR-4, a pattern-recognition receptor, promote pro-inflammatory cytokine secretion in response to stimuli such as Lipopolysaccharides (LPS) and Aβ aggregates^25,27^. We used TAK-242, a well-known TLR-4 inhibitor that has previously been shown to reduce pro-inflammatory cytokine secretion induced by Aβ aggregates^31^. We pre-treated iMGLs with TAK-242 for 20 minutes before adding *in vitro* or engineered Aβ aggregates and measured both Aβ internalization and cytokine levels post-exposure **(Fig. 3A)**. Our results show that TAK-242 effectively reduced uptake **(Fig. 3B-E)** and cytokine release, with notable decreases in TNF-α, IL-1β, and IL-6 levels for both *in vitro* and engineered Aβ42 **(Fig. 3F, H-J)** and Aβ40 **(Fig. 3 G, L-N)** aggregates. The impact of this inhibition (∼80%) was particularly significant in iMGLs treated with smaller 30nm aggregates **(Fig. 3K, O)**. However, the inhibitory effect was decreased substantially when using 100nm Aβ aggregates (∼40%) and was absent (∼0%) with larger 500nm aggregates for both Aβ42 and Aβ40. (**Fig 3. K, O**).

**Figure 3.**
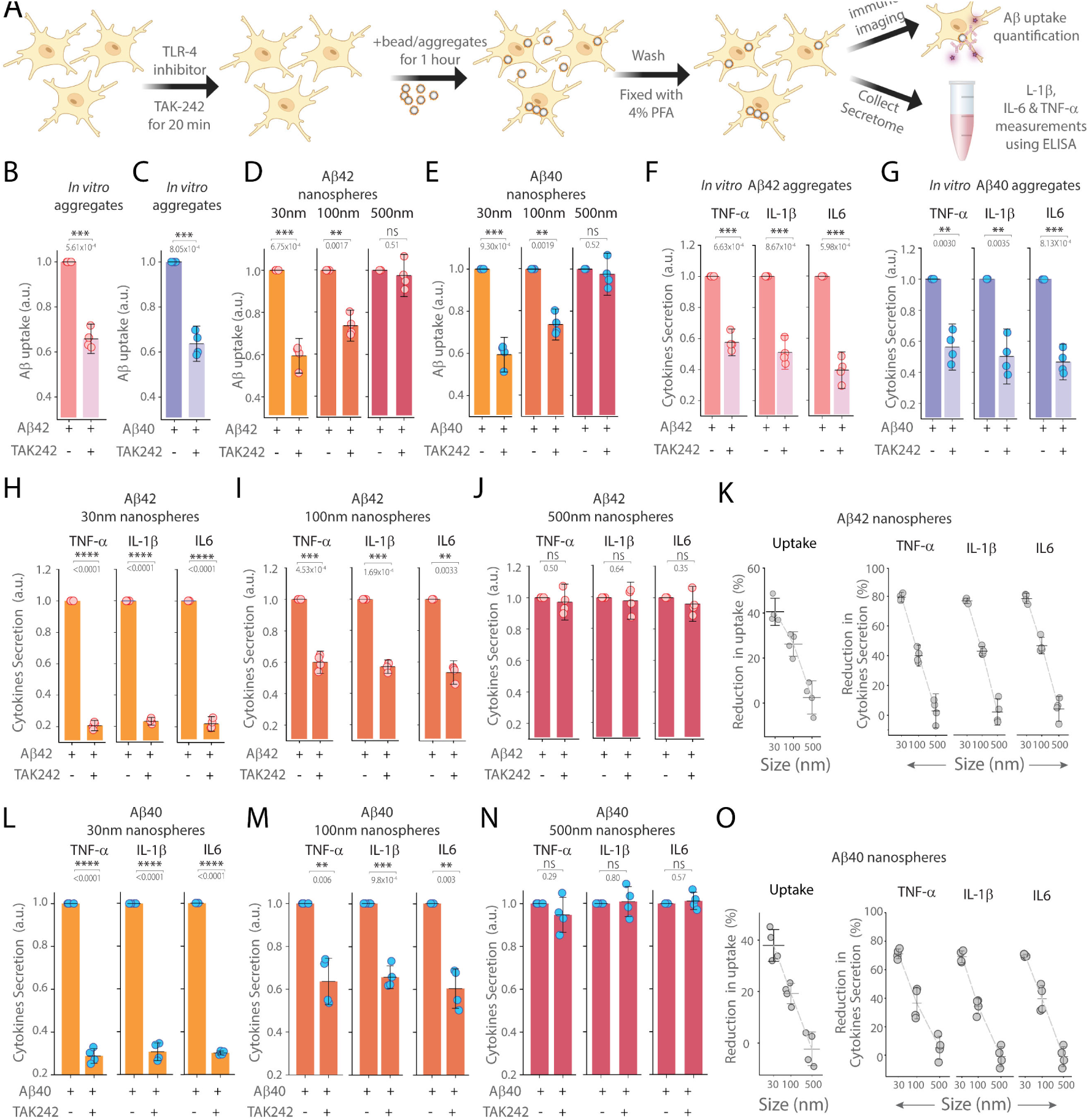
Effect of TLR4 inhibition by TAK-242 on iMGLs’ internalization of Aβ aggregates and cytokine secretion. **(A)** Schematic illustrating the protocol to assess Aβ uptake and cytokine levels using ELISA, with and without TLR-4 inhibitor TAK-242. **(B-C)** Quantification of Aβ internalization by iMGLs incubated with *in vitro* prepared aggregates of **(B)** Aβ42 and **(C)** Aβ40, comparing conditions with and without TAK-242. **(D-E)** Comparison of Aβ uptake by iMGLs incubated with **(D)** Aβ42 and **(E)** Aβ40 conjugated to nanospheres of various sizes (30nm, 100nm, 500nm), analyzed both with and without TAK242. For each case, internalization data are normalized to conditions without TAK-242. **(F-G)** Measurement of cytokine secretion (TNF-α, IL-1β, IL-6) by iMGLs following 1 hour incubation with *in vitro* prepared **(F)** Aβ42 and **(G)** Aβ40 aggregates, under conditions with and without TAK-242. **(H-O)** Measurement of cytokine release (TNF-α, IL-1β, IL-6) by iMGLs following 1 hour incubation with Aβ42 and Aβ40 aggregates conjugated nanospheres of different sizes: **(H, L)** 30nm, **(I, M)** 100nm, and **(J, N)** 500nm, with and without TAK-242. Reduction in cytokine secretion (TNF-α, IL-1β, IL-6) by iMGLs incubated with **(K)** Aβ42 and **(O)** Aβ40 conjugated nanospheres of different sizes (30nm, 100nm, 500nm) following TAK-242 treatment. For each case, cytokine release data are normalized to conditions where Aβ was added in the absence of TAK-242. Data are presented as the mean ± standard deviation across four biological replicates. Statistical significance was assessed using an unpaired two-sample t-test. *P < 0.05, **P < 0.01, ***P < 0.001, ns - non-significant (P ≥ 0.05)

### Subtle changes in protein-protein interactions shift protein aggregates’ functions from healthy to diseased states

Next, we explored how protein-protein interactions regulate the biological functions of protein aggregates. We focused on the interactions between Aβ40 and Aβ42, as their varying proportions significantly influence the rate of progression rate of AD^32–34^. To mimic the changes observed during the disease^32^, we studied the interaction of Aβ40 and Aβ42 in ratios of 9:1 and 7:3, alongside pure forms of each proteins. We aggregated Aβ40 and Aβ42 individually and in the specified ratios, measuring their aggregation kinetics using the ThT assay **(Fig. 4A)**. Our findings revealed that Aβ42 fibrillizes more rapidly, whereas Aβ40 aggregates at a slower pace, even when using concentrations ten times higher than Aβ42 to closely mirror their ratios in healthy conditions. When Aβ40 and Aβ42 were combined in the 9:1 and 7:3 ratios, the aggregation kinetics altered significantly compared to the pure forms and each other, indicating that Aβ40 and Aβ42 interact during the aggregation process in a manner dependent on their proportions. Notably, the 9:1 Aβ40:Aβ42 ratio aggregated slower than the 7:3 **(Fig. 4A)**, aligning with the trends observed in previous studies^35^.

**Figure 4.**
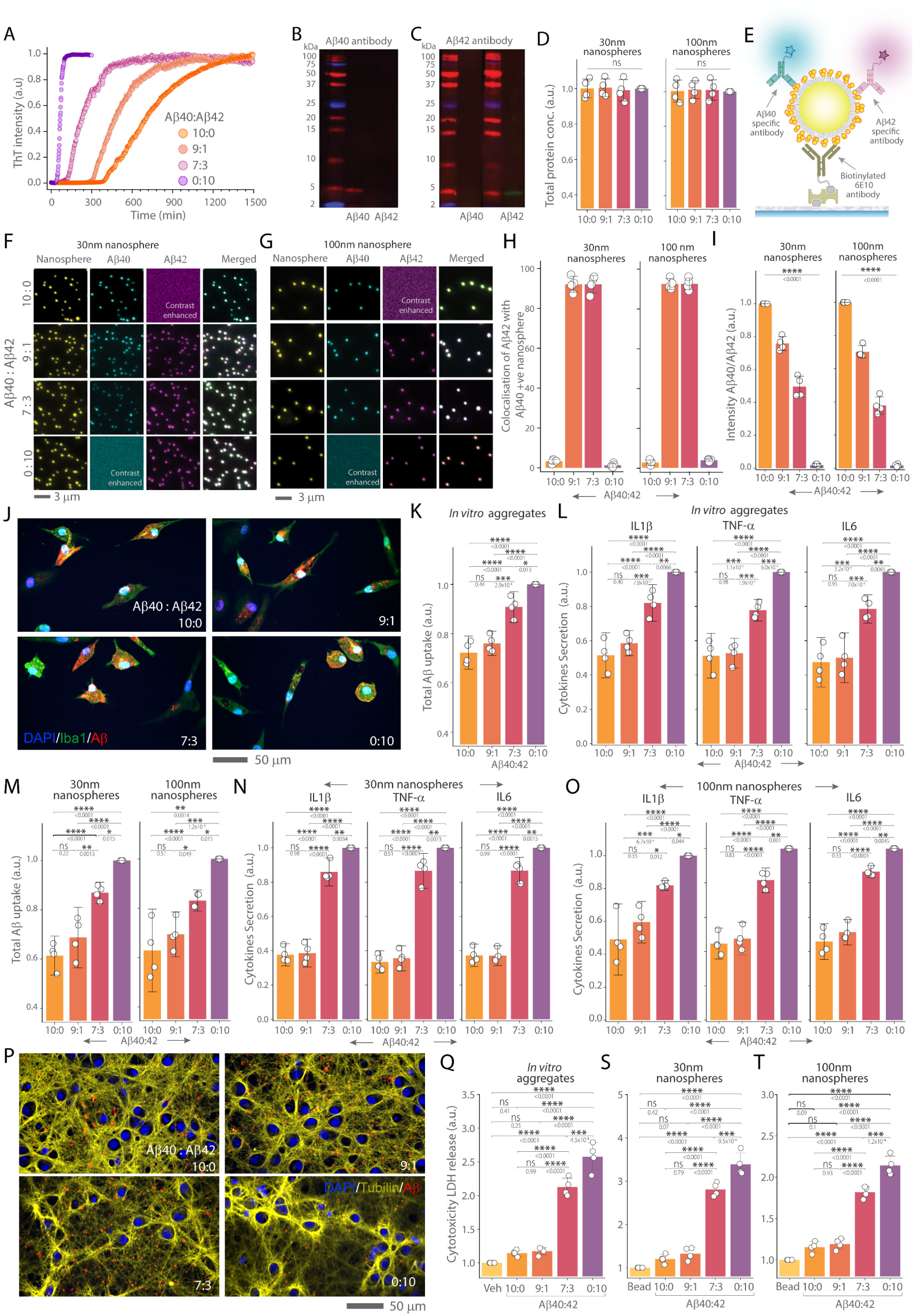
Effect of Aβ40 and Aβ42 ratio on the extent of iMGLs uptake, inflammatory cytokine secretion and neuronal toxicity. **(A)** ThT assay demonstrating the aggregation kinetics of Aβ40 (30µm), Aβ42 (3µm) alone, and in ratios of 9:1 (Aβ40 27µm, Aβ42 3µm) and 7:3 (Aβ40 21µm, Aβ42 9µm). **(B-C)** Western blot analysis of Aβ40 and Aβ42 using **(B)** Aβ40-specific EPR23712-2 antibody and **(C)** Aβ42-specific 21F12 antibody. **(D)** BCA assay to measure the total protein load attached to 30nm and 100nm nanospheres. For each nanosphere size, data are normalized to the Aβ42 aggregates engineered on the corresponding nanosphere. **(E)** Schematic representation of SiMPull analysis for engineered protein aggregated nanosphere. **(F-G)** Wide-field images of fluorescent nanospheres of 30nm and 100nm, conjugated with Aβ40 and Aβ42 alone and in ratios of 9:1 and 7:3. These aggregates are captured using 10nM biotinylated 6E10 antibody and imaged with 1nM Aβ40 specific and Aβ42 specific antibodies. Contrast-enhanced images are included to improve visualization where no signal from antibodies is observed. **(H)** Quantification of colocalization of Aβ40 coupled with Aβ42 on nanospheres at different Aβ40:Aβ42 ratios (10:0, 9:1, 7:3, 0:10). **(I)** Representative images of iMGLs (stained with DAPI/Iba1) showing internalization of Aβ aggregates at different Aβ40:Aβ42 ratios (10:0, 9:1, 7:3, 0:10). Aβ is stained with the 6E10 antibody. **(K)** Quantification of total Aβ internalization by iMGLs incubated with *in vitro* prepared aggregates of different Aβ40:Aβ42 ratios. Uptake data are normalised to pure Aβ42 aggregates. **(L)** Measurement of cytokine secretion (IL-1β, TNF-α, IL-6) by iMGLs after incubation with *in vitro* aggregates of different Aβ40:Aβ42 ratios. Cytokine release are normalized to the secretion levels in response to pure Aβ42 aggregates. **(M-O)** Quantification of total Aβ uptake and cytokine secretion (IL-1β, TNF-α, IL-6) by iMGLs following the incubation with 30nm **(M, N)** and 100nm **(M, O)** nanosphere-conjugated Aβ aggregates of different Aβ40:Aβ42 ratios. For both uptake and cytokine release, across both nanosphere sizes, data are normalized to pure Aβ42 aggregates. **(P)** Representative images of mouse primary neurons (stained with DAPI/acetylated Tubulin) treated with both *in vitro* and engineered aggregates with different Aβ40:Aβ42 ratios (10:0, 9:1, 7:3, 0:10). Aβ was detected with 6E10 antibody. **(Q-T)** Comparison of cytotoxicity, measuring LDH release from mouse primary neurons after exposure to *in vitro* prepared aggregates and, 30nm **(S)** and 100nm **(T)** nanospheres conjugated with different Aβ40:Aβ42 ratios. Data are normalized to values obtained when vehicle control of PBS buffers is used. Data points are plotted as the mean ± standard deviation representing three biological replicates. Statistical significance was calculated using one-way ANOVA with post-hoc Tukey mean comparison. *P < 0.05, **P < 0.01, ***P < 0.001, ns - non-significant (P ≥ 0.05).

To investigate how these proportions influence their disease-relevant functions, we co-aggregated Aβ40 and Aβ42 at the specified ratios (9:1 and 7:3) on 30nm and 100nm nanospheres, alongside their pure forms. We performed Western blot to choose a pair of antibodies specific to Aβ40 and Aβ42 that do not cross-react for further characterization **(Fig. 4B-C)**. We performed the BCA **(Fig. 4D)** on different sizes of conjugated nanospheres, finding no significant differences in protein load. Then the SiMPull assay was employed to further characterize these protein-conjugated nanospheres at the single-aggregate level **(Fig. 4E)**. These results showed Aβ40 and Aβ42 were colocalized over 90% when co-aggregated on the surface of nanospheres for both 9:1 and 7:3 ratios **(Fig. 4F-H)**. The the SiMPull analysis **(Fig. 4I)** and MSD assay (**SI Fig. 8**) confirmed that Aβ40 and Aβ42 are aggregated on the nanospheres in the intended ratios.

To assess how these changing ratios of Aβ isoform influence their biological functions, we utilized iMGLs **(Fig. 4J)** as previously described. We found that results from *in vitro* prepared aggregates, in terms of both uptake **(Fig. 4K)** and secretion of IL-1β, IL-6, and TNF-α **(Fig. 4L)**, agreed with those obtained from engineered aggregates **(Fig. 4M-O)**. Specifically, total Aβ uptake and secretion of pro-inflammatory cytokine increased as the proportion of Aβ40 decreased and Aβ42 increased within the co-aggregates.There was no difference in total Aβ internalization and cytokine release between pure Aβ40 aggregates and aggregates composed with a 9:1 Aβ40 to Aβ42 ratio, for both *in vitro* or engineered nanospheres **(Fig. 4K-O)**.

### Missense mutations at the same position differentially modulate protein functions

Next, we explored the impact of missense mutations on aggregates’ functions by focusing on the Dutch (E22Q) and Arctic (E22G) mutations which are associated with severe forms of AD due to their unique impact on Aβ42 aggregation and neurotoxicity^36,37^. These mutations are heterozygotic, leading to the expression of both mutant and WT Aβ isoforms in equal proportion^38^. Therefore, we performed the aggregation kinetics of each variant using ThT assay, both individually and in combination with WT Aβ42 at 1:1 molar ratio **(Fig. 5A-B)**. Our findings, similar to previously reported observations^39^, showed that the mutations significantly accelerated the aggregation of WT Aβ42. Co-aggregation rates falling between those of the mutant and WT Aβ42 alone, implying interactions between the mutant and WT during co-aggregation. To further investigate these interactions, we used 30nm nanospheres to prepare aggregates of WT Aβ42 alongside those with each mutation, both in pure forms (representing homozygosity) and in 1:1 molar ratio (representing heterozygosity).

**Figure 5.**
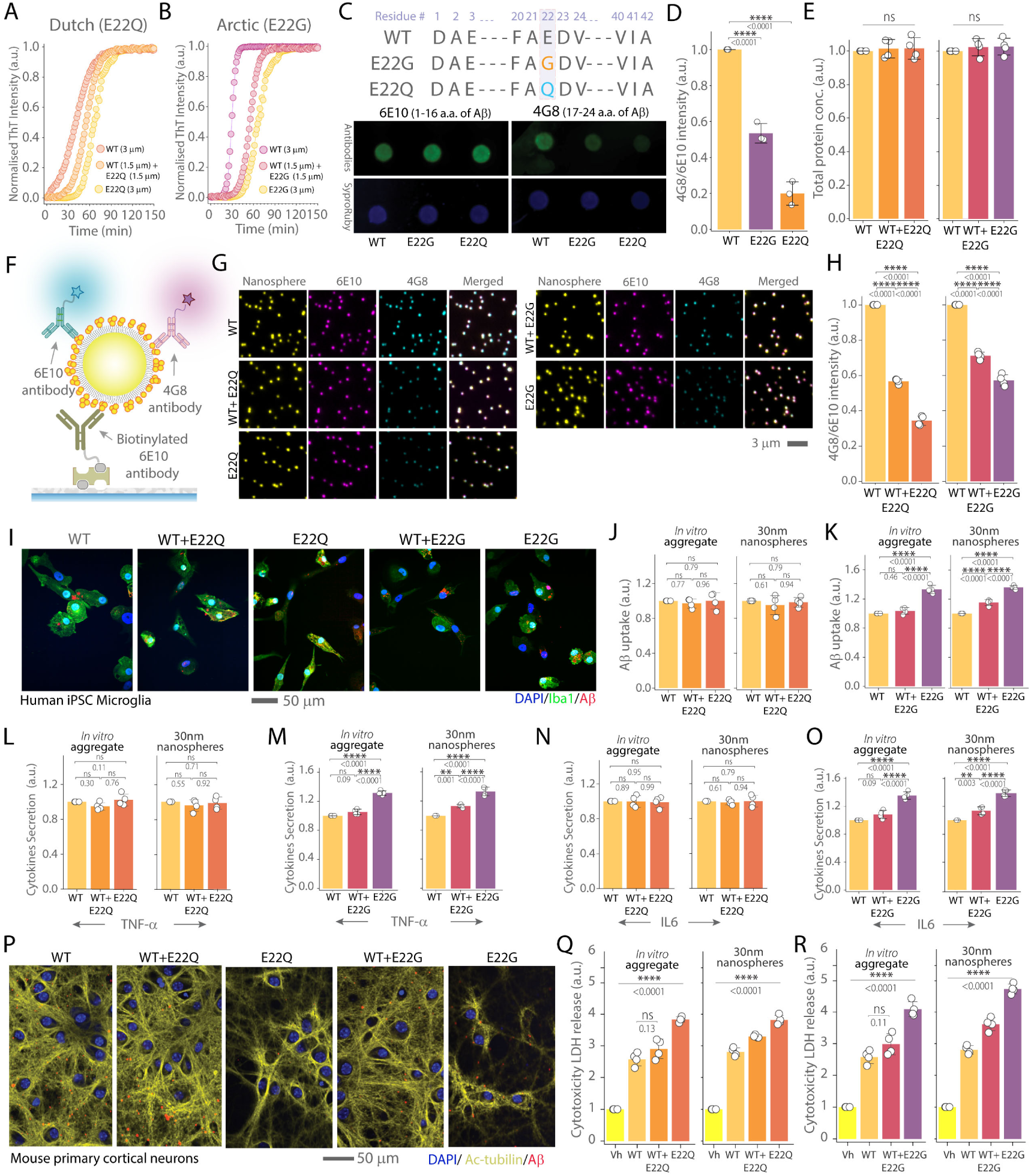
Impact of Dutch (E22Q) and Arctic (E22G) Aβ42 missense mutations on iMGLs uptake and neuronal toxicity. **(A-B)** ThT assay illustrating the aggregation kinetics of WT Aβ42, Dutch (E22Q) Aβ42, and Arctic (E22G) Aβ42 missense mutations both alone and in 1:1 ratios with WT Aβ42. **(C)** Illustration of amino acid change and dot blot analysis of WT, E22Q, and E22G using antibodies 6E10 and 4G8, with Sypro Ruby staining to estimate protein load on each well. **(D)** Quantification of the 4G8/6E10 intensity ratio for WT, E22Q, and E22G from dot blot experiments normalised to SyproRuby intensity. **(E)** BCA assay for total protein measurements for WT, E22Q, and E22G conjugated to 30nm nanospheres. **(F)** Schematic representation of SiMPull analysis for characterizing protein-conjugated nanospheres. **(G)** Wide-field fluorescence microscopy images displaying WT Aβ42, E22Q, WT + E22Q (1:1), E22G, and WT + E22G (1:1) aggregates conjugated to 30nm nanospheres and stained with 6E10 and 4G8 antibodies. **(H)** Quantification of 4G8/6E10 intensity ratio for each case. **(I)** Representative images of iMGLs (stained with DAPI/Iba1) showing internalization of different Aβ42 aggregates: WT, WT+E22Q, Aβ42 E22Q WT+E22G, and E22G. Aβ is stained with 6E10 antibody for quantification. **(J-K)** Quantification of total Aβ uptake by iMGLs incubated with *in vitro* prepared aggregates and 30nm nanosphere-conjugated Aβ42 aggregates for WT, WT + E22Q, and WT + E22G. Uptake data are normalized to corresponding WT Aβ42 uptake. **(L-O)** Measurement of cytokine secretion (TNF-α in **L, M** and IL-6 in **N, O**) by iMGLs following incubation with *in vitro* prepared aggregates and 30nm nanosphere-conjugated aggregates of WT, E22Q, E22G, WT + E22Q, and WT + E22G. Cytokine release data are normalized to responses from WT Aβ42. **(P)** Representative images of mouse primary cortical neurons (stained with DAPI/ acetylated tubulin) treated with WT, E22Q, E22G, WT+E22Q, and WT+E22G Aβ42 aggregates. **(Q-R)** Cytotoxicity assay measuring LDH release from mouse primary cortical neurons after exposure to *in vitro* prepared aggregates and 30nm nanospheres conjugated with different Aβ aggregates. Data are normalized to values from PBS buffer used as the vehicle control. Data presented as mean ± standard deviation across four biological replicates. Statistical significance was assessed via unpaired two-sample t-test (D) or one-way ANOVA with Tukey’s post-hoc test *P < 0.05, **P < 0.01, ***P < 0.001, ns non-significant (P ≥ 0.05).

To differentiate between mutant and wild-type forms of Aβ42, we faced a challenge due to the lack of mutation-specific antibodies. To circumvent this problem, we hypothesized that the 6E10 antibody, which targets the 1-16 residues of Aβ, would not distinguish between the mutant and wild-type forms. Conversely, we anticipated that the 4G8 antibody, targeting residues 17-24, would show reduced binding to the mutations due to alterations within this epitope. To test this hypothesis, we performed a dot blot analysis using 4G8 and 6E10 antibodies. The results confirm our expectations, showing a lower 4G8 to 6E10 intensity ratio for the mutant forms compared to the WT Aβ42 **(Fig. 5C-D)**. A BCA assay was used to measure total protein concentration, with no significant differences in total protein load observed among the WT and mutants individually, and the co-aggregates **(Fig. 5E)**. We then employed SiMPull analysis to further characterize the nanosphere-conjugated aggregates **(Fig. 5F).** Single-particle imaging and subsequent analysis corroborated the findings from the dot blot analysis, with both mutations demonstrating reduced 4G8 to 6E10 intensity ratios compared to the WT **(Fig. 5G-H)**. Homotypic aggregates of E22Q and E22G exhibited even lower 4G8 to 6E10 ratios than the corresponding mutant-WT Aβ42 co-aggregates as expected.

To explore how these Aβ42 mutations affect the microglial response, iMGLs were exposed to the WT, mutant and co-aggregates for 1 hour. We found differential microglial activation depending on the specific mutation. While the Dutch E22Q mutation did not significantly alter microglial uptake **(Fig. 5J)** or cytokine release **(Fig. 5L,N)**, the Arctic E22G mutation led to increased uptake **(Fig. 5K)** and enhanced IL-1β, IL-6, and TNF-α secretion compared to WT Aβ42 **(Fig. 5M,O)** from both *in vitro* and 30nm nanospheres. Finally, we assessed the neurotoxic potential of these mutations using mouse primary neuronal cultures. LDH assays revealed increased neurotoxicity when treated with the mutated Aβ42 compared to WT, with homotypic forms of E22Q and E22G aggregates showing higher toxicity than their respective WT-mutant co-aggregates **(Fig. 5Q-R)**. This differential toxicity between pure WT and WT-mutant co-aggregates was evident only in engineered aggregates, not in those produced by *in vitro* conventional methods. This distinction is likely due to the inherent heterogeneity of aggregates formed through conventional methods.

### The proportion of post-translational modifications dictates the damaging functions of protein aggregates

We then turned our focus to the role of pathological PTMs, which are widespread in neurodegenerative diseases^40,41^, to assess their impact on protein aggregate functions. Specifically, we explored into the phosphorylation of αSyn, a key event in PD pathogenesis^41^. We used immunohistochemical methods to examine fixed post-mortem midbrain tissues from three individuals with PD, identifying phosphorylated α-synuclein (pSyn) within Lewy bodies, a characteristic marker of PD^41–43^. Then, we extracted αSyn aggregates from the flash-frozen brain tissues from the same areas and employed SiMPull imaging. Our imaging results show significant but variable amount of colocalization of αSyn with pSyn.

To examine how this PTM influences their functions, we engineered nanospheres with varying ratios of αSyn to pSyn: 100:0, 75:25, 50:50, 25:75, and 0:100. We chose these ratios to mirror the conditions observed in healthy individuals and PD patients, where typically only 4% of α-Syn is phosphorylated in healthy individuals, whereas in Lewy bodies extracted from PD brains, over 90% of the α-Syn is phosphorylated^43^. Phosphorylation of αSyn was performed using the kinase PLK3, which specifically phosphorylates αSyn at S129^44^, and was characterized using Western blot **(Fig. 6D),** and SiMPull analysis **(Fig. 6E-F)**. BCA assay confirmed no significant difference in protein load across the different aggregate ratios **(Fig. 6G).** Then, we employed a biotinylated MJFR1 antibody to capture αSyn-containing aggregates for SiMPull analysis. Subsequently, we introduced two imaging antibodies, MJFR-14-6-4-2 and MJFR-phospho antibody, to detect αSyn aggregates and pSyn, respectively. Wide-field fluorescence imaging demonstrated over 90% of the fluorescent nanospheres were colocalised with both αSyn and pSyn for all engineered co-aggregates **(Fig. 6H)**.To verify the intended ratios at the single aggregate level, we analyzed the intensity ratios using antibodies specific to αSyn and pSyn, confirming that the proportion of pSyn increased within the aggregates led to a gradual increase in mean intensity ratio bteween pSyn and αSyn **(Fig. 6I).** Additionally, MSD assays verified that we successfully achieved the intended ratios of pSyn within the nanospheres **(SI Fig. 9).**

**Figure 6.**
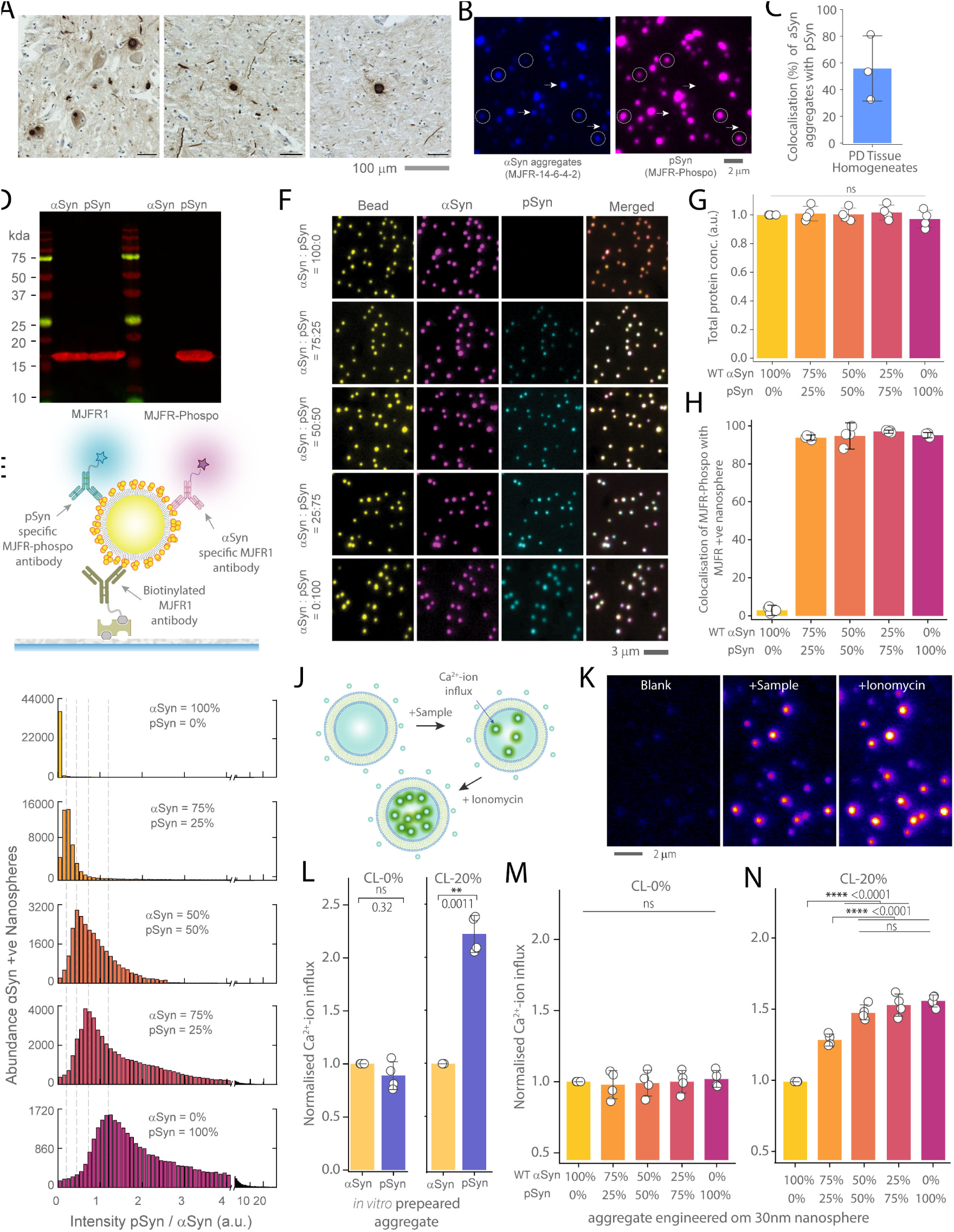
The proportion of phosphorylated αSyn within αSyn aggregates impacts its ability to permeabilize mitochondria mimicking membrane. **(A)** Immunohistology images displaying the presence of pathological pSyn-rich Lewy bodies and Lewy neurites in post-mortem midbrain tissue from three PD patients using pSyn/81A Ser129 phosphorylated antibody. **(B)** SiMPull images of αSyn aggregates and pSyn isolated from PD tissue homogenates. Dotted circles and arrows indicate colocalized and non-localized spots. **(C)** Quantification of colocalization levels of αSyn aggregates with pSyn in tissue homogenates. **(D)** Western blot analysis for αSyn and pSyn detection using MJFR1 and MJFR1-Phospho antibodies. **(E)** Schematic representation of the SiMPull analysis used for characterizing αSyn and pSyn-conjugated nanospheres. **(F)** Three-color epi-fluorescence images depicting various ratios of αSyn and pSyn (100:0, 75:25, 50:50, 25:75, and 0:100) engineered on 30nm fluorescent nanospheres (1µM monomer equivalents) using 10nM biotinylated Syn211 antibody for capture and αSyn confirmation specific Alexa-561-fluor conjugated 5nM MJFR-14-6-4-2 antibody in conjunction with pSyn-specific 5nM Alexa-637-fluor labelled Anti-Alpha-synuclein (phospho S129) antibody EP1536Y. **(G)** BCA assay to quantify protein load on nanospheres. Data are normalized to the WT αSyn aggregates on nanospheres. **(H)** Quantification of the three color colocalization of nanosphere, αSyn and pSyn at various αSyn and pSyn ratios. **(I)** The intensity ratios between pSyn and αSyn channels gradually increase as the proportion of pSyn within the αSyn aggregates rises. The intensity ratio of pSyn to αSyn is calculated by comparing the corresponding antibody intensities. **(J)** Schematic illustration of the experimental protocol for quantifying membrane permeabilization by measuring Ca^2+^-ion influx in response to aggregates. **(K)** Representative images show the Ca^2+^-ion influx into Cal-520 dye-filled that are sequentially treated with buffer, sample, and ionomycin. **(L-N)** Quantification of Ca^2+^-ion influx in individual vesicles, composed with and without cardiolipin, in response to aggregates composed of various αSyn and pSyn ratios. (**L**) Comparison of *in vitro* prepared aggregates using pure αSyn and pSyn. (**M-N**) Quantification of the Ca^2+^-ion influx into thousands of individual vesicles composed of 20% Cardiolipin **(N)** and without Cardiolipin **(M)**, responding to varying ratios of αSyn to pSyn (0%, 25%, 50%, 75%, 100%) engineered on the surface of 30nm nanospheres. All Ca^2+^-ion influx measurements are normalized to the response from WT αSyn aggregates. Data points represent the mean ± standard deviation from three independent replicates. Statistical significance was assessed using two-samples t-test (L) or one-way ANOVA followed by Tukey’s post-hoc mean comparison. *P < 0.05, **P < 0.01, ***P < 0.001, ns - non-significant (P ≥ 0.05).

To evaluate the disease-relevant functions of αSyn aggregates with varying proportions of pSyn, we utilized a quantitative assay that measures protein-induced membrane permeabilisation^16,45–47^. This assay assess membrane damage by quantifying calcium-ion influx into synthetic vesicles filled with a calcium-sensitive dye **(Fig. 6J,K)**. A higher influx of calcium-ion indicates increased membrane permeabilization, correlating with protein aggregates’ higher potential to disrupt membrane integrity. This method was previously applied to analyze αSyn aggregates’ toxicity, using oligomeric aggregates prepared using recombinant proteins^16^ and those isolated from tissue^45^, and stem cell models^46^. For this study, we created two types of vesicles: one simulating plasma membranes and another containing cardiolipin to mimic mitochondrial membranes. Cardiolipin, a phospholipid found almost exclusively in mitochondria, is crucial for mitochondrial bioenergetics and influences the pore-forming activity of αSyn oligomers in mitochondrial membranes, thereby driving neurodegeneration^48,49^. Our findings revealed that while there was no notable difference in membrane permeabilization across the different αSyn to pSyn ratios in cardiolipin-free vesicles, the inclusion of cardiolipin markedly increased the disruptive effects of pSyn-containing aggregates **(Fig. 6L-N)**. This enhancement was observed for both *in vitro* prepared aggregates **(Fig. 6L)** and those engineered on nanospheres **(Fig. 6M,N)**. Notably, aggregates with even just 25% pSyn caused significantly more membrane damage than pure αSyn aggregates in the presence of cardiolipin, with 50% pSyn aggregates showing damage comparable to pure pSyn aggregates.

## Discussion

Protein aggregation is a complex process that generates transient oligomeric intermediates of varying sizes and compositions^3–5^, often identified as the most toxic species in the aggregation pathway^4,10,11,22^. Commonly used top-down methods such as sucrose gradient fractionation and ultracentrifugation, or time point capture of the aggregation pathway are typically employed to enrich specific populations for functional characterization^7,15^ but struggle with challenges such as the lack of uniformity in size, composition, and aggregation state. Oligomeric aggregates are transient, particularly in the presence of monomers^12^, making analysis of how size and composition affect function complicated. Additionally, the frequent mutations and PTMs in these proteins add further complexity to their functional analysis. In response to these challenges, we have developed a robust method for the functional analysis of oligomeric aggregates, focusing on controlled preparation with specific sizes and compositions that closely emulate those found in human conditions.

We achieved controlled preparation by covalently attaching chosen proteins to nanospheres, followed by adding excess monomers to initiate controlled aggregation. This method ensures that aggregation occurs on the surface of the nanospheres, resembling the oligomers formed during the lag phase of protein aggregation. Simply coating the nanospheres without additional monomeric protein does not lead to aggregate formaion **(SI Fig. 10)**. Conversely, starting with an excess of protein without prior coating results in clumps due to the hydrophobic nature of the nanosphere surface promoting non-specific binding **(SI Fig. 11)**. Our method minimizes the formation of free aggregates not associated with nanospheres, as the surface significantly enhances the rate of aggregate formation^13,50^.

After establishing this platform, we explored how the size of Aβ40 and Aβ42 aggregates affects microglial uptake and the associated cytokine response. Microglial internalization of Aβ is a key pathway for the clearance of this protein in the brain^25,27^. Using human stem cell-derived iMGLs, we showed that the mechanisms contributing to the uptake and cytokine response to Aβ oligomers are highly size-dependent for both Aβ40 and Aβ42. In particular, the contribution of TLR-4 is more pronounced when smaller aggregates (30nm) interact with iMGLs than with larger aggregates (100nm) and it diminished substantially when the aggregate size increased to 500nm, **(Fig. 3K, O)**. These findings indicate that the physical dimensions and aggregation state of Aβ critically regulate microglial biological responses and their contribution to disease progression.

We also examined how protein-protein interactions determine protein aggregates’ function by focusing on Aβ40 and Aβ42, the predominant Aβ isoforms in the human central nervous system. The interplay between Aβ40 and Aβ42 influences their biological functions and impacts the progression of AD. The declining ratio of Aβ40 to Aβ42 in human biofluids correlates with amyloid accumulation in neural tissue^51^ and disease progression^33,34^. AD patients with Presenilin 1 gene mutations, who exhibit a higher Aβ40 to Aβ42 ratio, experience a later onset of the disease^52^. Studies in mice^53^ and *Drosophila melanogastor*^54^ have shown that elevated levels of Aβ40 can mitigate the toxicity induced by Aβ42 and extend lifespan, indicating a protective role for Aβ40. These protective effects are attributed to the differential aggregation rates of Aβ40 and Aβ42; Aβ40 aggregates more slowly and can slow down Aβ42 aggregation^55^. However, how changing ratios shift the biological functions of their co-aggregates from a healthy to a diseased state remains poorly understood. Our results indicate that increasing the Aβ40 content within co-aggregates leads to decreased pro-inflammatory cytokine secretion by iMGLs and reduced neuronal toxicity. Although a higher proportion of Aβ42 within the aggregates leads to more uptake by iMGLs, this increase clearance comes with a significant downside - a surge in the activation of chronic neuroinflammatory pathways. Typically, acute inflammatory stimuli like LPS or Aβ activate neuroprotective pathways that help to mitigate the temporary effects of these stimuli, thus encouraging the uptake of more Aβ42^29^. However, when these immune responses persist, they result in chronic inflammatory activation, which in turn increases Aβ production and aggregation, and activation of various downstream toxic pathways such as oxidative stress^26,29^. This positive feedback cycles of sustained pro-inflammatory signalling, increased Aβ aggregation and instigation of toxic pathways contributes to the neurodegeneration observed in AD^8,56^.

Our data revealed that increasing the ratio of Aβ40 can significantly mitigate the pro-inflammatory cytokine secretion and neuronal toxicity induced by Aβ42. At a physiological ratio of 9:1 (Aβ40 to Aβ42), co-aggregate-induced toxicity is similar to Aβ40. However, when the proportion of Aβ42 increases to 7:3, as often observed in AD, there is a corresponding rise in microglia-related inflammation and neuronal toxicity. Our innovative approach of preparing aggregates with controlled composition provides novel insights into the functions of these aggregates, insights not attainable with conventional methods. Our findings support the Aβ40’s role as a natural modulator, capable of counteracting the damaging effects of Aβ42 and its potential effectiveness of therapeutic strategies aimed at restoring the natural balance of Aβ isoforms.

We also investigated the impact of two missense mutations, Aβ42 Dutch (E22Q)^37^ and Aβ42 Arctic (E22G)^36^, associated with early-onset AD. both resulting from pathogenic mutations at the same position (E673) in the amyloid precursor protein. These mutations are inherited in an autosomal dominant pattern, leading to heterozygous expression that produces an equal proportion of WT and mutant Aβ^38^. Using 1:1 engineered co-aggregates with relatively uniform composition provided a distinct advantage over traditional methods that typically generate a heterogeneous mix of aggregates. This mix includes separate populations of pure WT, pure mutant, and WT-mutant co-aggregated forms, which obscures the specific effects and complicates the analysis. By engineering co-aggregate nanospheres, we found clear differences in cytokine release and neurotoxicity between WT Aβ42 and co-aggregated WT and mutants, which was obscured using conventional preparation of aggregates **(Fig 5J-O, Q-R)**.

Finally, we studied the role of PTM on the functions of protein aggregates, a feature often linked to various neurodegenerative diseases. We focused on the phosphorylation of αSyn (pSyn), a pathological hallmark in PD and a potential therapeutic target. pSyn interacts with cardiolipin - a lipid exclusive to mitochondria - disrupting mitochondrial membrane integrity and leading to dysfunction^41,43,46^. Given the critical role of αSyn phosphorylation in PD pathology, we systematically varied the levels of pSyn within αSyn aggregates and investigated the neurotoxic effects. We found that even a 25% presence of pSyn within αSyn aggregates significantly increases toxicity by disrupting lipid membranes enriched with cardiolipin. When half of the αSyn within the aggregates were phosphorylated, they showed toxicity levels comparable to those pure pSyn aggregates. In the absence of cardiolipin from the membrane, we did not observe the increased damaging effect of pSyn, reflecting its preferential interaction with this mitochondria-exclusive phospholipid. Our approach provides novel insights into how pSyn modulates the pathological behaviour of αSyn aggregates, enabling us to determine the specific amount of PTM needed to render the aggregates toxic to mitochondria, a level of detail not achievable with conventional aggregation methods.

Our approach to studying protein aggregates provides control over experimental conditions, yet it may not fully replicate natural aggregation processes in living organisms. The use of nanospheres could potentially modify the structure of protein aggregates, and their presence precludes the study of protein degradation. Thus, integrating this approach with conventional aggregation studies could significantly improve the effectiveness of the technique. Moreover, it is essential to use only freshly prepared samples for functional analysis to avoid protein clumping **(SI Fig. 12).**

In conclusion, our methodology overcomes longstanding challenges in analysing the structure-function relationships of transient protein aggregates. This approach enables us to elucidate the mechanistic pathways of four critical disease-associated events shaped by the functions of protein aggregates in proteinopathies. Our ability to control the size and composition of these aggregates has led to new understandings of how physical dimensions, protein-protein interactions, subtle compositional changes such as missense mutations, and PTMs function as molecular determinants of disease pathology. This work not only deepens our understanding of the molecular mechanisms underlying incurable human conditions like Alzheimer’s and Parkinson’s diseases but also lays a foundation for the functional characterization of protein aggregates across a broader spectrum of proteinopathies.

## Materials and Methods

### Ethics Statement

Ethical approval for the AD and PD post-mortem tissue was granted through the Sheffield Brain Tissue Bank, UK. For iMGLs, human skin fibroblast samples were obtained for a different study under the Yorkshire and Humber Research and Ethics Committee number: 16/YH/0155.

### Immunohistochemistry of post-mortem AD and PD tissue

Post-mortem brain tissue from individuals with AD and PD were provided by the Sheffield Brain Tissue Resource UK. Five**-** micron thick sections of formalin-fixed paraffin-embedded human tissue from AD and PD patients were used to perform immunohistochemistry for Phosphorylated α-Synuclein (pSyn) (Biolegend Cat. No. 825701) and Pan-Amyloid beta (Aβ) antibody 4G8 (Biolegend Cat. No. 800708) separately. Tissue was dewaxed and rehydrated using xylene and a series of alcohols, respectively. Endogenous peroxidase was blocked using 3% H_2_O_2_ in methanol for 10 minutes. Antigen retrieval was performed using a pH6 buffer in which samples were microwaved at 100% power for 10 minutes. After this, samples were placed in 100% formic acid for 3-5 minutes. To minimize non-specific binding, blocking was performed with 2.5% normal horse serum (VECTASTAIN Elite ABC-HRP Kit Mouse IgG, Vector Laboratories Cat. No. PK-6102) for 30 minutes. The primary antibody was incubated overnight at 4°C at a 1:500 dilution for pSyn and a 1:5000 dilution for Aβ. A biotinylated horse anti-mouse IgG secondary antibody was applied to the tissue for 30 minutes followed by an avidin-biotin-peroxidase complex (VECTASTAIN Elite ABC-HRP Kit Mouse IgG, Vector Laboratories Cat. No. PK-6102) for 30 minutes. To visualise the antibody, 3,3’-diaminobenzidine (DAB) was used as the chromogen and hydrogen peroxidase as the substrate (DAB Substrate Kit, Vector Laboratories Cat. No. SK-4100) and incubated for 2-3 mins. Sections were counterstained with haematoxylin, dehydrated through a series of alcohols and mounted using dibutylphthalate polystyrene xylene (DPX). Scanned images of the staining were taken on the Hamamatsu NanoZoomer XR Slide scanner at 20X magnification. Cropped images were obtained on QuPath and scale bars were added in ImageJ.

### Extraction of diffusible aggregates from AD post-mortem tissue

Soluble Aβ aggregates were extracted from human tissue samples using a published protocol^17^. Tissue samples used in this study are summarized in supplementary table 1. Initially, the tissues from frontal cortex were sectioned into 200 mg chunks. These sections were then treated with 1 mL of artificial cerebrospinal fluid which composed of 120mM NaCl (Merck Cat. No. S9888), 2.5 mM KCl (Merck Cat. No. P3911), 1.5 mM NaH_2_PO_4_ (Merck Cat. No. S5011), 26mM NaHCO_3_ (Merck Cat. No. S6297), 1.3 mM MgCl_2_ (Merck Cat. No. M8266), at pH 7.4 and incubated at 4°C for 30 minutes under mild agitation. After this, the samples were centrifuged at 2,000 g for 10 minutes at 4°C. About 80% of the resulting supernatant was transferred and centrifuged again at 14,000 g for 110 minutes at 4°C. The upper 80% of the supernatant from this second spin was then dialyzed against a 50-fold excess of fresh aCSF buffer for 72 hours at 4°C, using a 2kDa molecular-weight-cut-off Slide-A-Lyzer dialysis cassette (ThermoFisher Cat. No. 66203); the buffer was refreshed every 12 hours. Post-dialysis, the samples were aliquoted, frozen at −80°C, and used for further experiments.

### Extraction of aggregates from PD post-mortem tissue

Post-mortem brain tissue of PD patients were homogenized using a previously published protocol^45^. Post-mortem tissue samples used in this study is tabulated in supplementary table 1. Midbrain tissue weighing 200 mg were homogenized in 1 mL of tris-buffered saline (20 mM Tris HCl, 500 mM NaCl, pH 7.5) containing protease inhibitor cocktails (Merck, Cat. No. 11697498001). The homogenates were first centrifuged for 5 minutes at 1000 × g at 4°C to remove highly insoluble debris. The resulting supernatants were then centrifuged for 30 minutes at 175,000 × g. The supernatant was collected for the characterization.

### Aggregation of recombinant Aβ peptide

The monomeric recombinant peptide was prepared by resuspending it in a solution of 1% NH_4_OH (Merck Cat. No. 221228) in PBS buffer at a concentration of 1 mg/mL, following the manufacturer’s instructions. To remove any insoluble components, the mixture was centrifuged at ∼4,000 g for 30 seconds. Next, the peptide solution was diluted into PBS at a concentration of 200 or 100 µM and 50-µL aliquots prepared on ice. The aliquots were then flash-frozen on liquid nitrogen and stored at −80 °C for future use. Subsequently, these aliquots were diluted in PBS to a total concentration of 3µM and aggregated in a 96-well half-area plate (Corning, Cat. No. 3881) at 37°C without shaking. The aggregation process was monitored using 20µM Thioflavin T dye (Sigma, Cat. No. T3516) using a plate reader (Clariostar Plus, BMG Biotech).

To analyze the species formed during Aβ42 aggregation, samples of the aggregation mixture (aggregated without ThT) were taken at the end of the lag phase or at the plateau phase. The time points used for analysis as well as detail of the peptide concentration and vendors are presented in the Supplementary table 2.

### dSTORM protocol and data analysis

We performed the direct stochastic optical reconstruction microscopy (dSTORM) was performed using the SiMPull method. Biotinylated 6E10 antibodies were used to capture Aβ aggregates, and Alexa-fluor 647 labeled 6E10 antibodies were used for the imaging. After the imaging antibody incubation of the SiMPull method, the PBS buffer was removed and then 100 mM MEA in Tris buffer (Idylle lab) was added as an oxygen scavenging system. This solution was freshly prepared immediately before imaging. Then with an exposure time of 30ms a total of 3,000 frames per acquisition were used using 647 nm laser illumination. The positions of the “blinking” events in the dSTORM images were determined using the Peak Fit module of the GDSC plugin Single Molecule Light Microscopy plugin package for ImageJ. The analysis performed using a signal strength threshold of 40 (a.u.) and a precision threshold of 30nm, with a magnification of eight. Finally, the sizes of individual aggregates were estimated using Gaussian fitting.

### Antibody conjugation

To prepare antibodies for SiMPull assays, we used the Lightning-Link conjugation kit from Abcam to attach biotin or Alexa Fluor dyes (488, 568, 637) to unlabeled antibodies according to the manufacturer’s instructions. First, we added 1µL of modifier reagent to 10µL of the antibody solution, followed by gentle mixing. This antibody-modifier mixture was then combined with the lyophilized conjugation mix and incubated for an hour. Post-incubation, 1µL of quencher was added to stop the reaction, mixed gently, and allowed to sit for 5 minutes. The conjugated antibodies were then stored at 4°C for subsequent experimental use.

### Phosphorylation of αSyn

Phosphorylation of αSyn was carried out according to a published protocol^44^. We used the polo like family of kinases, PLK3, which specifically phosphorylates αSyn at S129 more than 95%. Phosphorylation buffer (50mM HEPES, 1 mM MgCl_2_, 1mM EGTA, 1mM DTT) was freshly prepared and combined with 300µg αSyn along with 2mM Mg-ATP and 1µL PLK3. The solution was thoroughly mixed by pipetting and incubated at 30°C for 12 hours without agitation. Phosphorylation of αSyn at S129 (pSyn) was confirmed by western blotting using MJF-R13 alpha-synuclein phospho S129 antibody (Abcam Cat No. ab168381).

### Nanosphere coupling procedure

**F**or our experiments, we utilized two types of nanospheres -fluorescent nanospheres (ThermoFisher Cat No. F8888) for SiMPull imaging, and non-fluorescent nanospheres (ThermoFisher Cat No. C37486, C37269, C37274) for cellular and membrane permeation experiments, MSD assay, BCA assay. The size of these nanospheres were 30nm, 100nm, and 500nm. We procured the smallest nanospheres, advertised as 20 nm; however, upon TEM analysis (SI Fig 1), we discovered their mean diameter was 28.9 nm. Throughout the manuscript, we refer to these as 30 nm nanospheres.

We began by preparing 100mL of 50mM MES buffer at pH 6.0, dissolving MES sodium salt (Sigma Cat. No. M3885-25G) in Milli-Q water. The nanospheres resuspended in the buffer through gentle pipetting and were sonicated for 30 minutes to disrupt clumping. The protein coupling reaction was initiated by thawing aliquots of proteins (either WT Aβ42, WTAβ40, Aβ mutants, αSyn, or pSyn) intended for conjugation. These proteins were diluted to a concentration of 5 μM in MES buffer, and the calculated volume was added to each reaction mixture containing the resuspended nanospheres, with all components kept on ice. To ensure uniform reaction conditions across all nanosphere sizes, we calculated the surface area of the differently sized nanospheres to maintain a constant total surface area across samples. The amount of protein required was determined to ensure complete coverage of the nanospheres’ surface area, considering that the diameters of Aβ and αSyn are 1nm^20,57^, respectively. Following a 15-minute incubation on ice, EDC solution was added to each reaction mixture in MES buffer to achieve a final concentration of 1 mg/mL. The pH of each solution was then adjusted to 6.5. The reactions were incubated overnight at 4°C. To quench the reaction, a glycine solution was added to reach a final concentration of 0.1M. The mixture was then sonicated on ice cold water for 15minutes. The reaction mixtures were transferred to Lo-Bind Eppendorf tubes and centrifuged at 10,000 g, followed by three PBS washes. Between each wash a 15-minute sonication on ice was performed. The final protein-conjugated nanospheres were resuspended in PBS and stored at 4°C. Then the nanospheres were sonicated on ice for 30 minutes, then resuspended in PBS and supplemented with five times more monomers than the calculated surface area required, allowing aggregate formation overnight under quiescent condition at 4°C. The prepared aggregates were stored in the dark at 4°C and used within 36 hours for further experiments. For experiments involving mixtures of proteins, the intended ratios were used both for covalent coupling and during the aggregation process.

### BCA assay

To determine total protein load engineered on nanosphere surface, the Pierce BCA Protein Assay (Pierce Cat. No. 23225) was used. First, BSA standards were prepared in PBS to generate a standard curve, and samples were diluted to an appropriate concentration (1:10). 25 µL of standard or sample was applied to each well, in triplicate, in a clear 96-well microplate, and 200 µL reagents (BCA reagent A and B at 50:1) were added to each well. The plate was placed on a shaker for 30 seconds to induce mixing, and then incubated at 37°C for 30 minutes. The microplate was then cooled to room temperature and absorbance was measured using the Clariostar microplate reader.

### Wide-field fluorescence Imaging setup

We performed imaging using a custom-built microscope based on a Nikon Eclipse Ti2 body equipped with a Perfect Focus unit and three Omicron Luxx lasers (488 nm, 561 nm, and 635 nm). Lasers were launched through a fibre coupler (KineFlex SM/PM Fiber), collimated with Zoom Fiber Collimators (Thorlabs Cat No. C20FC-A), and passed through achromatic Quarter-Wave Plates (Thorlabs Cat. No. AQWP05M-600). Then the laser directed into the back focal plane of a 100x 1.49 NA oil-immersion objective lens (Nikon). To achieve uniform illumination, we integrated a beam shaper (Asphericon, Cat No. TSM25-10-LD-D-532) in the excitation path, resulting in less than 5% intensity variation across the imaging field **(SI Figure 13).** Fluorescence emissions were collected through the same objective and separated using a dichroic beamsplitter (Laser2000 Cat No. Di01-R405/488/561/635). The light then passed through a specific set of optical filters for each fluorophore before being captured by a Photometrics Prime 95B sCMOS camera. For each fluorophore, a combination of long-pass and band-pass filters were used: a 488 nm long-pass (Laser2000 Cat No. BLP01-488R-25) and a 530/50 nm bandpass (Laser2000 Cat No. FF01-530/55-25) for Alexa Fluor 488; a 561 nm long-pass (Laser2000 Cat No. LF561/LP-C-000) and a 593/46 nm bandpass (Laser2000 Cat No. FF01-593/46-25) for Alexa Fluor 561; and a 647 nm long-pass (Laser2000 Cat No. BLP01-647R-25) and a 680/42 nm bandpass (Laser2000 Cat No. FF01-680/42-25) for Alexa Fluor 647. The setup was controlled by Micro-Manager 2.0. The data acquisition was performed using automatic stage movements ensuring unbiased data collection. Images were averaged over 50frames at 50ms exposure each. For SiMPull imaging of nanospheres and super-resolution imaging of brain derived aggregates, we used epifluorescence and total internal reflection fluorescence modes, respectively.

### Single-molecule Pull down method

In SiMPull method, we followed a previously published protocol^4^. For Aβ aggregates, we utilized 10 nm of Biotinylated 6E10 (Biolegend, Cat. No. 803007) as the capture antibody. For the imaging probes, we employed a selected combination of Alexa-Fluor-647-labeled 6E10 (Biolegend, Cat. No. 803021), Alexa-Fluor 594 labeled 4G8 (Biolegend, Cat. No. 800716), Alexa-Fluor 561 labeled A11 Polyclonal Antibody (ThermoFisher, Cat. No. AHB0052), Alexa-Fluor 561 labeled Anti-beta amyloid 1-40 EPR23712-2 (AbCam, Cat. No. ab289991), and Alexa-Fluor 674 labeled Anti-Amyloid Beta [21F12] (Absolute Antibodies, Cat. No. Ab02391-3.0) or Amytracker 540 (Ebba biotech). All imaging antibodies were used at a concentration of 1nM and Amytracker 540 used at a concentration of 20nM in PBS. For αSyn aggregates, 10nM of Biotinylated Syn211 (Abcam, Cat. No. ab206675) was used as capture antibody and 5nM of Alexa-Fluor 561 labeled pSyn specific antibody and 5nM of Alexa-Fluor 637 labeled Anti-αSyn aggregate antibody [MJFR-14-6-4-2] (Abcam, Cat. No. ab214033) served as detection probes. Information on the capture and imaging probes for each experimental figure is tabulated in Supplementary Table 3. Glass coverslips (VWR, Cat. No. MENZBC026076AC40) were PEGylated and stored in a desiccator at −20°C to ensure cleanliness and functionality. We prepared the assay wells by initially coating them with 0.2mg/mL NeutrAvidin (ThermoFisher, Cat. No. 31000) in PBS containing 0.05% Tween-20 for 5 minutes. Following this, the wells were washed twice with 10µL of PBS containing 0.05% Tween-20 and once with 10µL of PBS containing 1% Tween-20. We then added 10µL of the appropriate biotinylated capture antibody in PBS containing 0.1mg/mL BSA (ThermoFisher, Cat. No. 10829410) to each well and allowed it to incubate for 15minutes. After the incubation period, the wells were washed following the same protocol as before. We then added the sample, which could be either in vitro prepared and engineered aggregates or brain extracts and incubated it for 1hour. Following another series of washes, a mixture of imaging probes in PBS containing 0.1 mg/mL BSA was introduced to the wells and incubated for 30minutes. The final washing step was performed twice with 10µL of PBS containing 0.05% Tween-20 and once with 10µL of 1× PBS containing 1% Tween-20. To finalize the preparation, 3μL of PBS was added to each well, and the samples were sealed with a second plasma-cleaned coverslip to ensure a controlled environment for subsequent imaging.

### Analysis of colocalisation data

The averaged images acquired with excitation at 488 nm, 561 nm and 635 and analysed using Fiji plugin ComDet3^58^. For spots in two/three different channels to be considered colocalized, the displacement between their centres of mass (determined by Gaussian fitting) was required to be ≤3 pixels. Colocalization by aggregate number was defined as the ratio between the number of colocalized spots and the total number of spots in the specific channel.

### Atomic force microscopy (AFM) and scattering-type, scanning near-field optical microscopy (s-SNOM)

Simultaneous AFM and s-SNOM scans were performed using a neaSCOPE from Attocube systems AG/Neaspec. The AFM was performed in tapping mode, with Pt/Ir coated ARROW-NCPt cantilevers from Nanoworld, at a tapping frequency of 289 kHz and a tapping amplitude of 89-91 nm. AFM height and phase maps were recorded, with the height maps corrected for sample tilt with a planar gradient offset. For the collection of s-SNOM data, light from a broadband illumination source (TOPTICA Photonics, FemtoFiber dichro midIR) with output approximately in the range 900-2000 cm^-1^ was sent into a Michaelson interferometer. One arm of the interferometer housed the AFM in operation on the sample, and the other arm housed a clean reference mirror. Light focussed onto the metal coating of the AFM cantilever tip (radius around 25 nm) generated surface excitations with a strong near-field component. These near-field electromagnetic fields interacted with the sample, generating a scattering centre. Further incoming light scattered off this interaction region between the cantilever and the sample, and was collected back through the interferometer to be interfered with the clean reference light. Lock-in detection was used to demodulate the signal at the second harmonic of the tapping frequency, in order to reduce background interference. The amplitude of this signal was plotted after being normalized to the maximum recorded value. We note that due to the diverse spectral nature of the illumination source, only limited background removal and limited optical characterization can be performed. However, due to the strong difference in optical properties between the sample (Aβ42 protein, polystyrene nanospheres) and the substrate (silicon), strong contrast can be observed in the data that allows for the identification of the Aβ42-shell on the surface of the nanospheres.

### Transmission electron microscopy

Samples were prepared by adding 5µL of samples onto glow-discharged, carbon-coated copper grids. Each sample was allowed to absorb for 1minute before blotting and washing twice with distilled water and once with 0.75% uranyl formate. It was then stained with 0.75% uranyl formate for 20 seconds, blotted to remove excess stain, and vacuum dried. The prepared grids were examined using a Tecnai Spirit T12 Microscope at 80 kV. Images were captured on a bottom-mount CCD camera with magnifications ranging from 1,200 to 68,000x and underfocus between 500-3000 nm.

### Single-molecule FRET measurements

Monomeric solutions of Aβ42 labeled at the N-terminus with HiLyte Fluor 647 (AnnSpec Cat. No. AS-64161) and HiLyte Fluor 488 (AnnSpec Cat. No. AS-60479-01) were reconstituted in 10 mM NaOH. The protein concentration was determined using the absorbance of HiLyte Fluor 647 (250,000 M^−1^ cm^-1^) and HiLyte Fluor 488 (70,000 M^−1^ cm^−1^). Then monomeric Aβ42 were flash-frozen after aliquoting and kept at −80 °C until further use. For *in vitro* aggregation, aliquots of labelled monomeric Aβ42 were diluted to an equimolar concentration of 1.5µM of HiLyte647 Aβ42 and HiLyte488 Aβ42 in PBS and incubated at 37 °C aggregation under quiescent conditions. For coupling with nanosphere, we use equal amount labelled Aβ42. The samples were added to the poly-L-lysine coated coverslips and incubated for 20 minutes and then washed twice using PBS. Then the FRET assay were performed using epi-fluorescence mode using 488 nm excitation and the emission was collected using a combination of a 647 nm long-pass and a 680/42 nm bandpass filters.

### Dot blot assay

We loaded 750 ng of Aβ42 peptide onto a 0.2 µm nitrocellulose membrane of the Bio-Dot Apparatus (Bio-Rad Cat. No. 1706545). Then the membrane washed twice with TBS containing 2% formaldehyde and then incubated with the same solution for 30 minutes. Then the membrane was rinsed with deionized water thrice. fixed with 7% MeCOOH and 10% MeOH for 15 minutes and stained with SYPRO™ Ruby Protein Blot Stain (Invitrogen, Cat. No. S11791) for 15 minutes. Post-staining, the membrane was washed thrice in deionized water, dried, and imaged using an Odyssey XF Imager (LI-COR) at 600 nm. It was further processed with a 10-minute wash in 150 mM Tris, pH 8.8, 20% methanol, rinsed, and air-dried. We then blocked the membrane with 5% low IgG BSA in TBS (Serva, Cat. No.11948) for 1 hour at room temperature, followed by overnight incubation at 4°C with 6E10 (BioLegend, Cat. No. 803002) or 4G8 (BioLegend, Cat. No.800701) antibodies at 1:8000. After three washes in TBS, secondary antibodies Alexa Fluor 680 Anti-Rabbit IgG (Jackson ImmunoResearch Laboratories Cat. No. 711-625-152) and Alexa Fluor 790 Anti-Mouse IgG (Jackson ImmunoResearch Laboratories Cat. No. 715-655-150) were applied at a 1:50,000 dilution in TBS. Following a 1-hour room temperature incubation and three TBS washes, the membrane was imaged with the Odyssey XF Imager.

### Western blot assay

We began by mixing 100 ng of Aβ40 or Aβ42 peptides in PBS with 2x Tris-Tricine sample buffer (Bio-Rad, Cat. No. 1610739) and loaded them onto a 16.5% Mini-PROTEAN Tris-Tricine Gel (Bio-Rad, Cat. No. 4563063) or a 16.5% Criterion Tris-Tricine Gel (Bio-Rad, Cat. No. 3450064). We also loaded 8 µL of Precision Plus Protein Dual Xtra Prestained Protein Standards (Bio-Rad, Cat. No. 1610377). The samples were resolved in Tris-Tricine running buffer (Bio-Rad, Cat. No. 1610744) at 90V at 4°C until the dye reached the gel’s bottom. We then transferred the proteins to a 0.2 µm nitrocellulose membrane (Amersham, Cat. No. GE10600001) using a Criterion blotter (Bio-Rad, Cat. No. 1704070) in Towbin transfer buffer at 100V for 20 minutes. The membrane was incubated in 2% FA in PBS for 1hour with 5% milk in TBS-T for another hour at room temperature, and then incubated overnight at 4°C with primary antibodies: Anti-beta Amyloid 1-40 [EPR23712-2] (AbCam, Cat. No. ab289991) for Aβ40 and Anti-Amyloid Beta [21F12] (Absolute Antibodies, Cat. No. Ab02391-3.0) for Aβ42. After three 10-minute washes with TBS-T, we applied secondary antibodies Alexa Fluor 680 Anti-Rabbit IgG (Jackson ImmunoResearch Laboratories, Cat. No. 711-625-152) and Alexa Fluor 790 Anti-Mouse IgG (Jackson ImmunoResearch Laboratories, Cat. No. 715-655-150) at a 1:50,000 dilution for 1hour. Following three more washes, we visualized the proteins using an Odyssey XF Imager.

### Human iPSC-derived microglia-like (iMGL) Culture and immunocytochemistry

Induced pluripotent stem cells (iPSCs) from a 52-year-old female donor were cultured using mTeSR+ medium (StemCell Technologies, Cat. No. 85850) on vitronectin-coated plates until about 80% confluence, then passaged with Relesr (StemCell Technologies, Cat. No. 100-0484) and seeded on Matrigel-coated plates at 140,000 cells per 10cm². For differentiation, iPSCs were exposed to E8 media (StemCell Technologies, Cat. No. 05990) supplemented with 1% penicillin-streptomycin (ThermoFisher, Cat. No.15140122), 10 µM Rho kinase inhibitor (ROCKi) (StemCell Technologies, Cat. No. 72304), 5ng/ml BMP4 (Peprotech, Cat. No. 120-05ET), 1 µM CHIR99021 (Axon, Cat. No. 1386), and 25ng/ml activin A (Peprotech, Cat. No. 120-14P) at 37°C in a 5% O_2_ and 5% CO_2_ atmosphere. After 24 hours, media was changed to the same but with 1 µM ROCKi. At 44hours, cells transitioned to FVI media containing DF3S media (DMEM/F12 (ThermoFisher, Cat. No. 11320033), GlutaMAX (ThermoFisher, Cat. No. 35050038), 0.5% penicillin-streptomycin (ThermoFisher, Cat. No. 15140122), L-ascorbic acid (Sigma, Cat. No. A4403), Na_2_SeO_3_ (Sigma Cat. No. S5261), NaHCO_3_ (Sigma, Cat. No. S6014) supplemented with FGF2 (Peprotech, Cat. No. 100-18B), SB431542 (Stemcell technologies, Cat. No. 72232), insulin (Sigma, Cat. No. I9278), and VEGF (Peprotech, Cat. No. 100-20). After an additional 24 hours, cells shifted to normoxic conditions and cultured in HPC media DF3S base with FGF2, insulin, VEGF, TPO (StemCell Technologies, Cat. No. 78210), SCF (StemCell Technologies, Cat. No. 78155), IL-6 (StemCell Technologies, Cat. No. 78148), and IL3 (StemCell Technologies, Cat. No. 78146), changed daily for four days until cobblestone cell patches appeared. Progenitor cells were then collected, filtered, and seeded in ultra-low attachment dishes (Corning Cat. No.16855831) with Proliferation media (IMDM (ThermoFisher, Cat. No. 12440053), FBS (ThermoFisher, Cat. No. 16000044), insulin, MCSF (StemCell Technologies, Cat. No. 78150), and IL-34 (StemCell Technologies, Cat. No. 100-0930). For uptake experiments, iMGLs were incubated with 1µm monomer equivalent Aβ aggregates, then washed and fixed and stained for imaging. For TLR-4 inhibition, cells are treated with TAK-242 for 20 minutes before the addition of Aβ aggregates. Antibody Anti-Iba1 (Abcam, Cat. No. ab178846) and 6E10 (Biolegend, Cat. No. 803001) are used for immunocytochemistry, followed by secondary antibodies (ThermoFisher, Cat. Nos. A-11029 and A-31573) and DAPI staining (ThermoFisher, Cat. No. 62248). Cells were imaged with an Opera Phenix High Content Imaging System (PerkinElmer).

### Data analysis of cellular uptake assay

Data analysis was performed using Harmony High-Content Imaging and Analysis Software. For each set, images from ten randomly selected fields per well were captured across three wells. The imaging employed three channels: 405 nm for nuclear staining, 488nm for cell boundary visualization using Iba1 in iMGLs, and 647 nm for detecting Aβ with an Alexa Fluor-647 labeled 6E10 antibody. We delineated cell boundaries for iMGLs based on their respective staining. Masks generated from the Iba1 channels separated cellular from non-cellular areas. These masks were then used on the Aβ channel to measure fluorescence intensities, facilitating the quantification of Aβ uptake. Background fluorescence, derived from cells stained only with secondary antibodies, was subtracted to ensure measurement accuracy.

### Meso Scale Discovery assay

We measured Aβ levels using the V-PLEX Plus Aβ Peptide Panel 1 (6E10) Kit (MSD, Cat. No. K15200E), following the manufacturer’s guidelines. Initially, plates were blocked with Blocker A (MSD) for one hour at room temperature. Samples were then added and incubated for one hour. Subsequently, SULFO-TAG-labeled anti-human Aβ 6E10 antibodies were added to the plates and incubated for another hour. Throughout all incubation phases, the plates were agitated on an orbital shaker at 800 rpm. After completing three wash cycles, Read Buffer (MSD) was applied to the plates. The resultant signals were detected using a MESO QuickPlex SQ 120 multiplexing imager. For αSyn detection, a similar protocol to the one used for Aβ was followed. MFR1 (Abcam Cat no. ab138501) per well was used as the capture antibody, and an anti-human synuclein antibody from MSD along with anti-alpha-synuclein phospho S129 antibody EP1536Y (Abcam Cat. no. ab209422) were used as imaging antibodies for total and pSyn, respectively.

### ELISA to measure cytokine and chemokine concentrations in cell media

To measure cytokine and chemokine secretion by iMGL cells, the cell media were collected after a 1-hour incubation with various aggregates and then stored at −80°C for later analysis. The concentrations of IL-1β, IL-6, and TNF-α in the cell media were determined using the Duoset® enzyme-linked immunosorbent assay (ELISA) development systems (R&D Systems, Cat. No DY201, DY206, DY210), following the manufacturer’s instructions.

### Primary neuronal culture

First culture plates were coated with 200µL of poly-D-lysine (0.1mg/mL in dH_2_O, Merck-Sigma, Cat. No. P6407) and incubated overnight at 37°C in a 5% CO_2_ environment. Time-mated C57BL/6 female mice were ordered from Charles River, UK, and kept at the University of Sheffield Biological Services Unit until they were ready to be sacrificed by cervical dislocation. Cerebral cortices were isolated from embryonic day 15 embryos while submerged in cold HBSS−/− (Gibco, Cat. No. 14170088). Meninges were manually removed, and cortices were dissected using surgical forceps. The tissue was washed once in 10mL of HBSS−/−, then resuspended in 5 mL of HBSS−/− containing 0.05% trypsin (Gibco, Cat. No. 15090046) and incubated for 15 minutes at 37°C. After incubation, 5 mL of HBSS+/+ (Gibco, Cat. No. 24020117) supplemented with 10 μg/mL DNAse (Merck, Cat. No. 04536282001) was added for 2minutes, and the supernatant was then aspirated. The tissue was resuspended in 1mL of triturating solution composed of 1% Albumax – (Gibco Cat. No. 11020021), 0.5mg/mL trypsin inhibitor (Merck Cat. No. T9003), 10μg/mL (DNAse in HBSS−/−) and triturated through flame-polished glass Pasteur pipettes (ThermoFisher Cat. No. 11765098) with progressively smaller openings to obtain a single cell suspension. Cells were then resuspended in Neurobasal Plus media (Gibco Cat. No. A3582901) supplemented with B27 Plus supplement (Gibco Cat. No. A3582801), 2mM GlutaMax (Gibco Cat. No. 35050061), and 50unit/mL Penicillin/Streptomycin (Gibco Cat. No. 15070063) and maintained at 37°C in a 5% CO_2_ environment. Half media changes were performed every 3-4 days until the cultures reached 14 days. For neurotoxicity measurements, cells were incubated with Aβ aggregates (equivalent to 3 µm monomer) for 6hours and then fixed and stained for imaging. Antibody incubation used Anti-acetylated tubulin (ThermoFisher, Cat. No. 32-2700) and 6E10 (Biolegend, Cat. No. 803001), followed by secondary antibodies (ThermoFisher, Cat. Nos. A-11029 and A-31573) and nuclear staining Hoechst 33342 (ThermoFisher, Cat. No. 62249). Cells were imaged with an Opera Phenix High Content Imaging System.

### LDH cytotoxicity assay

LDH Cytotoxicity Assay Kit (ThermoFisher, Cat. No. C20303) were used to measure toxicity in primary neuron culture, we added in vitro prepared or engineered aggregates at a concentration of 1µM (monomer equivalent) on the 14th day’s culture and incubated them for 6 hours. After incubation, the cells were washed three times with PBS. We then collected the cell supernatant to assay for lactate dehydrogenase (LDH) activity. As a positive control, we used the supernatant from cells treated with RIPA lysis buffer (ThermoFisher, Cat. No. 89900), and as a negative control, we used the medium from untreated neurons. We added 100µL of the reaction mixture provided by the kit for detecting LDH activity following the manufacturer’s instructions. We stopped the reactions after 30 minutes using the stop buffer and measured the absorbance at 480 nm using a Clariostar plus plate reader (BMG Biotech).

### Membrane permeabilization assay

Membrane permeabilization assays were performed using a previously published method^47^. The lipid composition for the vesicles was used to mimic the mitochondrial membrane^48^ which included 30% 16:0–18:1 PC (Avanti Lipids, Cat. No. 850457), 40% 16:0-18:1 PE (Avanti Lipids, Cat. No. 850757), 20% 18:1 Cardiolipin (Avanti Lipids, Cat. No. 710335), 3% 16:0 SM (Avanti Lipids, Cat. No. 860584), 3% 16:0-18:1 PI (Avanti Lipids, Cat. No. 850142), 3% 16:0-18:1 PS (Avanti Lipids, Cat. No. 840034), and 1% biotinylated 18:1-12:0 Biotin PC (Avanti Lipids, Cat. No. 860563C). Control membranes were prepared without cardiolipin. Vesicles with 200 nm mean diameter were prepared by extrusion and ten freeze-thaw cycles, hydrated in 100 μM Cal-520 dye (Stratech, Cat. No. 21141) in 50 mM HEPES buffer of pH 6.5 and immobilized on argon plasma (Deiner Zepto One) cleaned coverslips (VWR, Cat. No. 6310122). Coverslips are the coated with PLL-g-PEG (20kDa PLL grafted with 2kDa PEG and 3.5 Lys units/PEG Chains, SuSoS AG) and PLL-g-PEG biotin (20 kDa PLL grafted with 2 kDa PEG and 50% 3.4kDa PEG-Biotin, SuSoS AG) in 100: 1 ratio at ∼1 mg/mL. Then 50µL of 0.1mg/mL NeutrAvidin (ThermoFisher, Cat. No. 31000) in HEPES buffer was added to the coverslips. Then the vesicles were immobilised on the coverslip and imaged with Ca^2+^-containing buffer (ThermoFisher, Cat. No. 21083027) (F_blank_), then exposed to the samples for 15 minutes and reimaged (F_sample_), this was followed by ionomycin treatment (Cambridge Bioscience, Cat. No. 1565-5) for Ca^2+^ -ion saturation (F_ionomycin)_. Relative Ca^2+^ influx was calculated using: = (F_sample_ - F_blank_)/ (F_ionomycin_ - F_blank_). Fluorescence emission of Cal-520 dye were passed through filters (BLP01-488R-25 and FF01-520/44-25, Laser 2000) before being imaged using a Photometrics Prime 95B sCMOS camera. Images were acquired at a power density of ∼10 Wcm^-2^ with a scan speed of 20Hz.

## Data availability

All data are available from the corresponding authors upon request. Source data are provided in this manuscript.

## Author Contributions

A.U., E.E.P and A.O. synthesized engineered the aggregates. A.U., E.F.G., E.E.P., N.G., and E.L.B did the protein aggregation and SiMPull experiments. M.C.K, E.E.P., and H.E.W. performed the cellular uptake assay. E.E.P. modified antibodies. A.O. and A.K. performed the AFM-NSOM experiments. E.F.G. performed the histopathology and. A.U. conducted neuronal toxicity measurements. A.U. and E.E.P. performed western and dot blot experiments and measured cytokines in media. E.E.P. did the membrane permeabilization and A.U. performed LDH assays. J.R.H. provided the post-mortem tissue samples. Data analysis, interpretation and statistics were conducted by A.U., E.F.G., E.E.P and S.D. This study was supervised by A.I.T., T.M., J.R.H. and S.D. The initial draft of the paper was written by A.U., E.F.G., and S.D.; all other authors critiqued the output, provided feedback, and contributed to editing the manuscript into its final form. S.D. conceived and designed the study. All authors read and approved the manuscript.

## Acknowledgments

We are grateful to the Sheffield Brain Tissue Bank for supplying the tissue and to those who have donated tissue for scientific research and their families who have supported this. This study was supported by the UKRI Future Leaders Fellowship (MR/V023861/1) (E.F.G., A.U., and S.D.), an EPSRC grant 2594676 (H.E.W, S.D) and Academy of medical sciences springboard award (SBF006\1038) (S.D.). The Sheffield NIHR Biomedical Research Centre provided support for this study.

