## Supporting info file for "Molecular determinants of protein pathogenicity at the single-aggregate level"

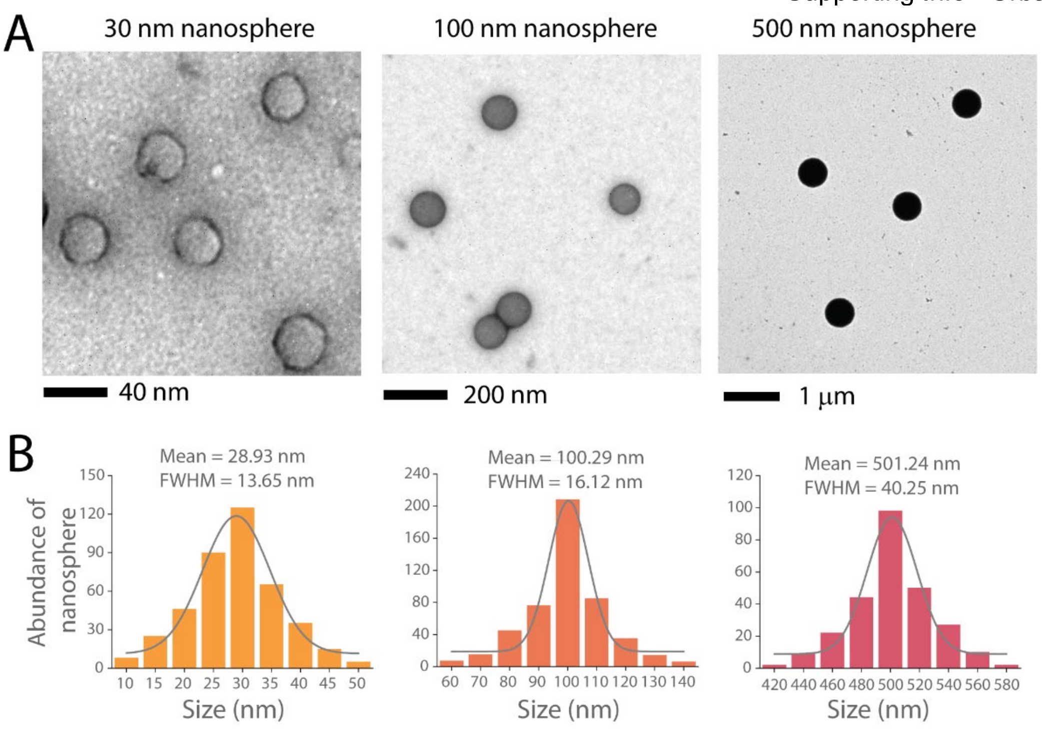


**Supplementary Figure 1. (A)** Representative TEM images of nanospheres in three different size ranges. **(B)** Size distribution for each size range of nanosphere. The smallest nanospheres, initially specified as 20nm by the vendors, have a measured mean diameter of 28.93nm and a full width at half maximum of 13.65 nm; these will be referred to as 30nm nanospheres throughout the manuscript. The other size ranges include nanospheres with mean sizes of 100.29nm and 501.24nm, respectively, labelled as 100nm and 500nm nanospheres.

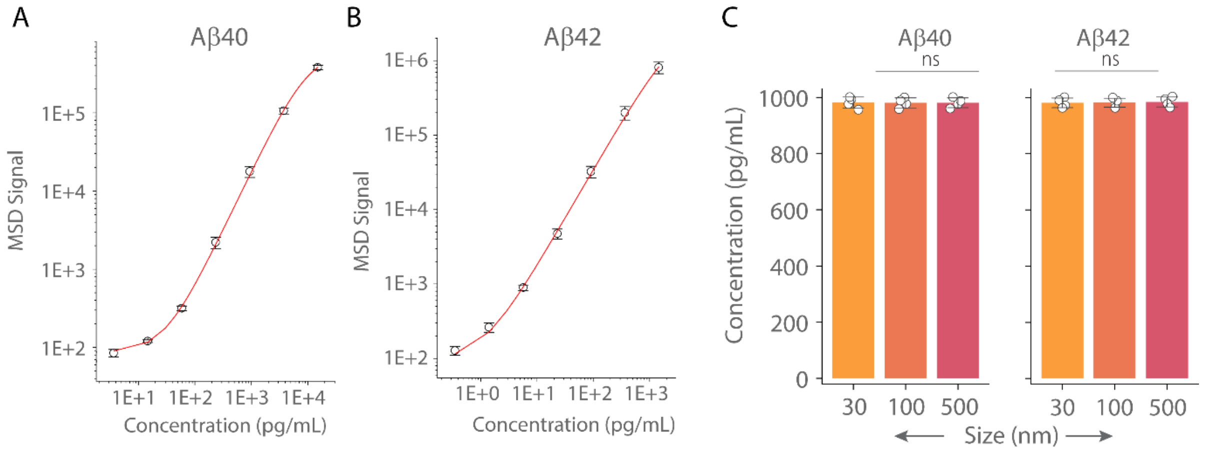


**Supplementary Figure 2.** Standard curve for **(A)** Aβ40 and **(B)** Aβ42 detection using the MSD assay, performed with the V-PLEX Aβ Peptide Panel 1 (6E10) Kit. **(C)** MSD assay results quantifying the protein engineered to 30nm, 100nm, and 500nm nanospheres. Data are presented as the mean ± standard deviation across four biological replicates. Statistical significance assessed using an unpaired two-sample t-test. *P < 0.05, **P < 0.01, ***P < 0.001, ns - non-significant (P ≥ 0.05).

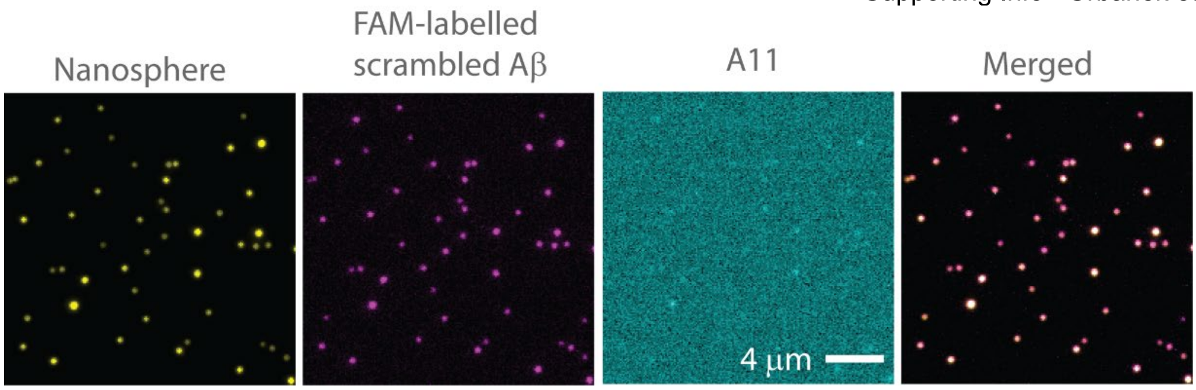


**Supplementary Figure 3.** FAM-labelled scrambled Aβ42 covalently conjugated and then placed on the surface of 30nm nanospheres. For this, we utilized 'dark-red' (Em680) carboxy-modified nanospheres instead of the yellow-green (Em520) ones used predominantly in the study, to prevent overlap with FAM (Em –520). These species are immobilized on a poly-L-lysine coated surface and imaged using wide-field imaging. High colocalization between scrambled Aβ42 and the nanospheres confirms that the proteins are conjugated to the nanospheres, but the lack of Alexa Fluor-561 labelled A11 antibody binding indicates that scrambled Aβ42 did not form aggregates on the surface as expected.

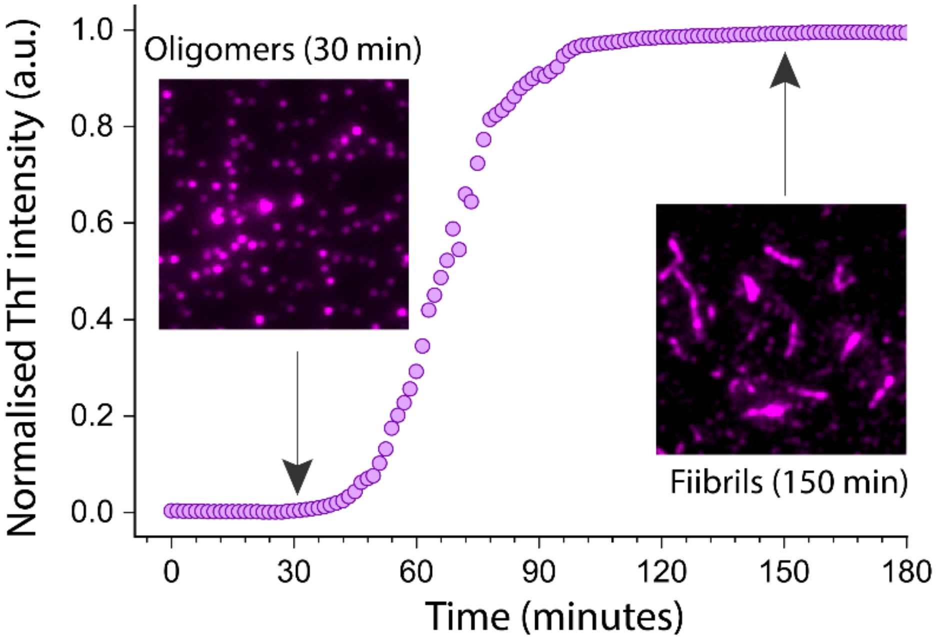


**Supplementary Figure 4.** Aggregation kinetics of Aβ42 monitored using the ThT assay. Aggregates formed at 30 and 150 minutes of aggregation reaction are isolated and characterized using the SiMPull assay, with biotinylated 6E10 antibody used for capture and Alexa Fluor 647 labelled 6E10 antibody utilised for imaging. At the lag phase of aggregation (30 minutes), the aggregates primarily consist of oligomers, while at the plateau phase (150 minutes), the aggregates are predominantly in fibrillar form. These samples are utilized in the FRET assay as shown in Figure 1O.

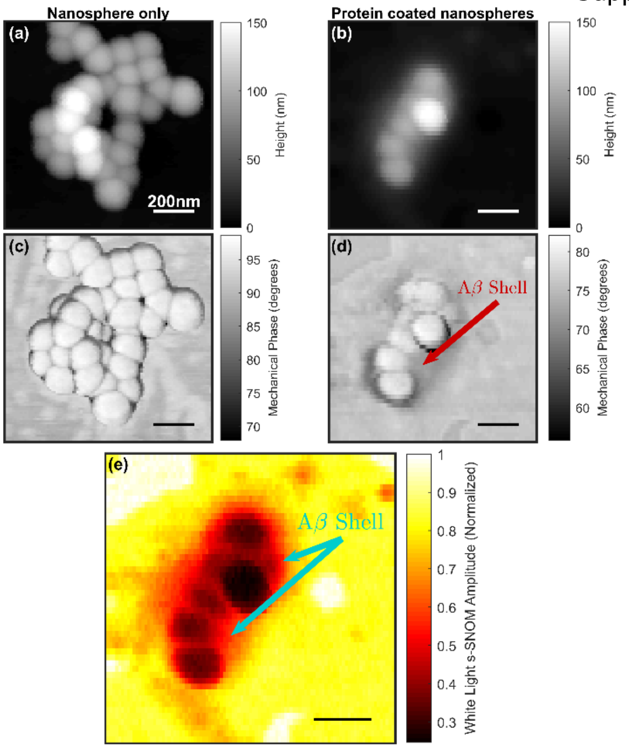


**Supplementary Figure 5.** Atomic force microscopy (AFM) and scattering-type, scanning near-field optical microscopy (s-SNOM) images of uncoated and Aβ42-coated 100nm nanospheres. **(a)** and **(b)**: AFM topography scans taken from the uncoated and coated Nanospheres, respectively. **(c)** and **(d)** AFM phase data pertaining to the scans shown in **(a)** and **(b)** respectively. AFM phase data can relate to the mechanical properties of the sample, with an area of (d) highlighted as an area of contrast that relates to the Aβ42-shell on the surface of the nanospheres. **(e)** Normalized s-SNOM amplitude data pertaining to the AFM data in **(b)** and **(d)**. Illumination was provided with a broadband source in the mid-infrared, with output approximately from 900 $cm^{-1}$ to 2000$cm^{-1}$.

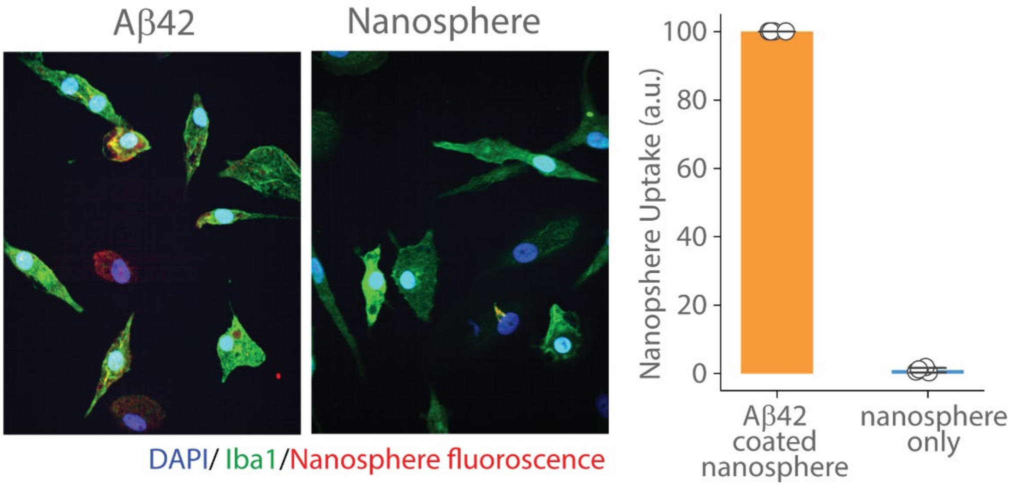


**Supplementary Figure 6.** Representative images of iMGLs (stained with DAPI/Iba1) following exposure to engineered Aβ42 aggregates on 30nm nanospheres and nanospheres alone (uncoated). Uptake was quantified using the intrinsic fluorescence of the nanospheres and normalised with engineered Aβ42 aggregates.

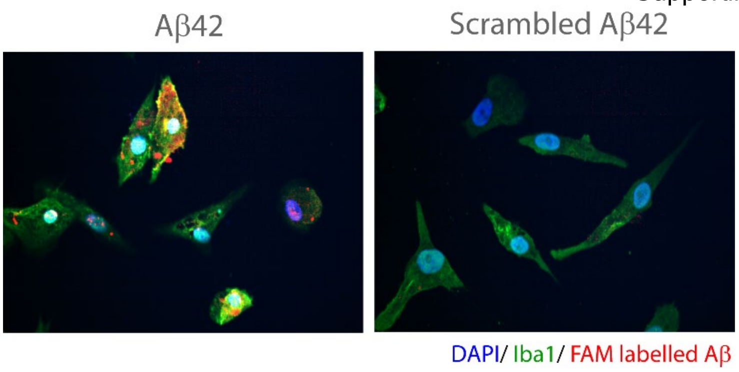


**Supplementary Figure 7.** Representative images of iMGLs (stained with DAPI/Iba1) following the uptake of FAM-labelled Aβ42 and FAM-labelled scrambled Aβ42 aggregates conjugated to 30nm nanospheres. Instead of using immunohistochemistry as done throughout the manuscript, we quantified the uptake with dye-labelled proteins due to the lack of a specific antibody for scrambled Aβ42.

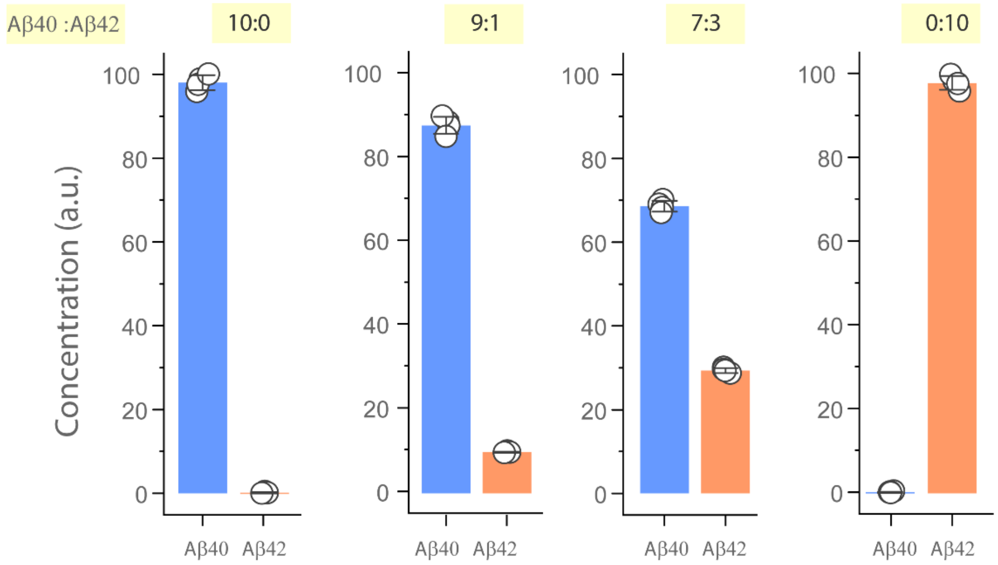


**Supplementary Figure 8.** MSD assay quantification of Aβ40 and Aβ42 engineered on the surface of 100nm nanospheres at different ratios (9:1 and 7:3), alongside pure forms of each protein. This data also demonstrates that the antibodies used to detect Aβ42 and Aβ40 do not cross-react.

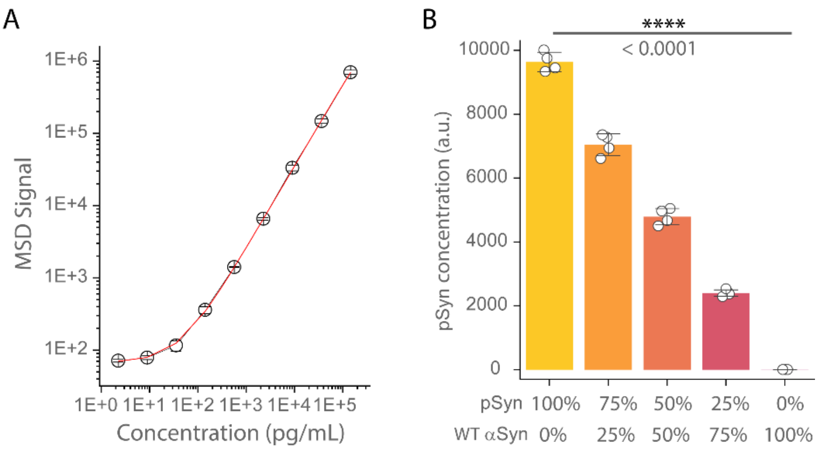

**Supplementary Figure 9. (A)** Standard curve for pSyn detection using the MSD assay, employing a biotinylated MJFR1 antibody for capture and a pSyn-specific Alexa-637-fluor labeled Anti-Alpha-synuclein (phospho S129) antibody EP1536Y for detection. **(B)** MSD measurements of pSyn concentration within αSyn aggregates (ratios of 100:0, 75:25, 50:50, 25:75, and 0:100) engineered on 30nm fluorescent nanospheres (1μM monomer equivalents). Data are presented as the mean ± standard deviation from four biological replicates. Statistical significance assessed using an unpaired two-sample t-test. *P < 0.05, **P < 0.01, ***P < 0.001, ns - non-significant (P ≥ 0.05).

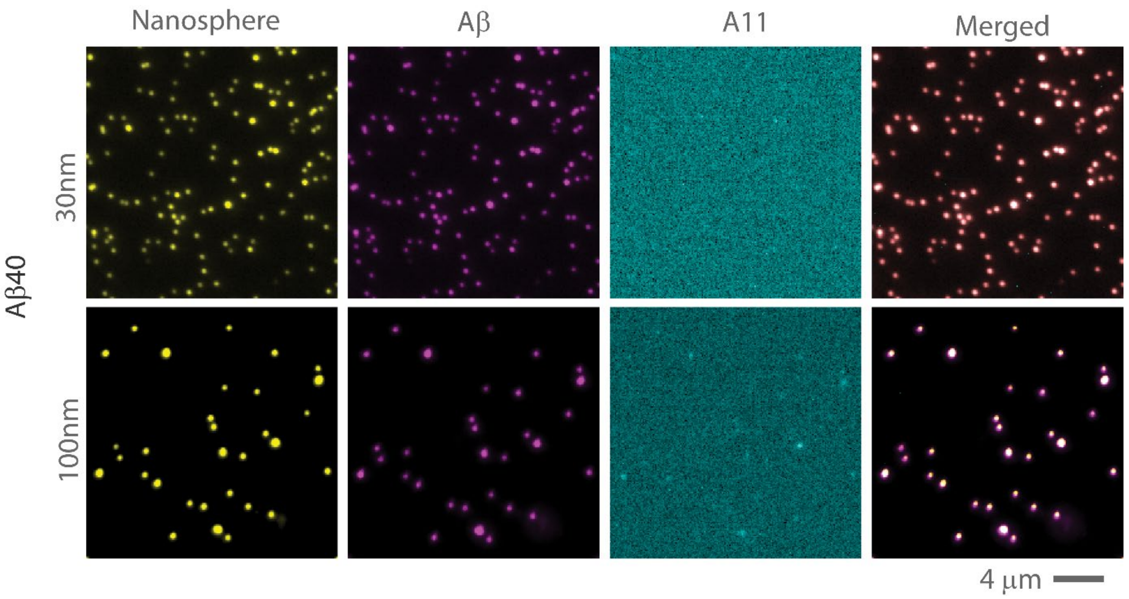


**Supplementary Figure 10.** SiMPull images of 30nm and 100nm nanospheres covalently conjugated with Aβ42, without the second step of aggregation using excess monomer. For bother sizes, Aβ conjugated nanospheres bind to the Aβ-specific 6E10 antibody, but the oligomer-specific A11 antibody does not bind to them. This demonstrates that simply coupling Aβ on nanospheres does not result in aggregate formation.

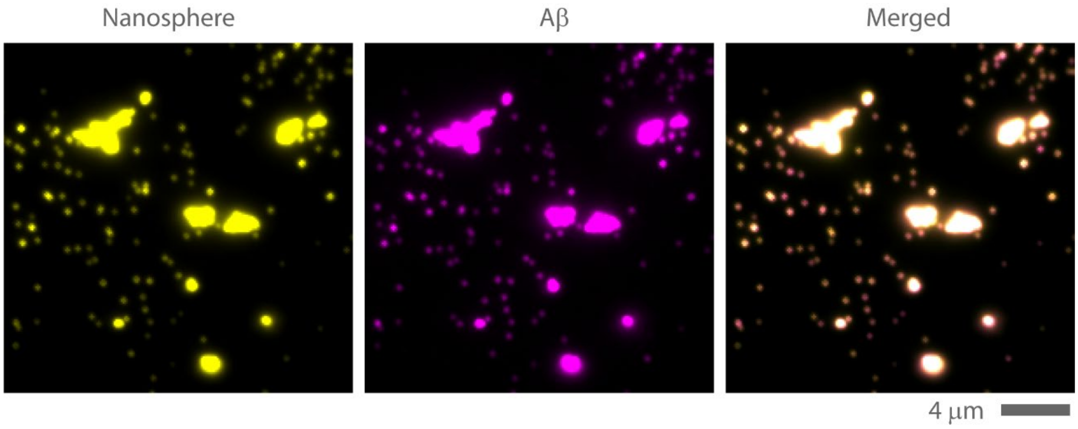


**Supplementary Figure 11.** SiMPull images show Aβ42 aggregates engineered on the surfaces of 30nm nanospheres without the initial covalent conjugation step. The Aβ42 aggregation to the nanospheres in the absence of initial coating leads to the clumping of nanospheres and aggregates.

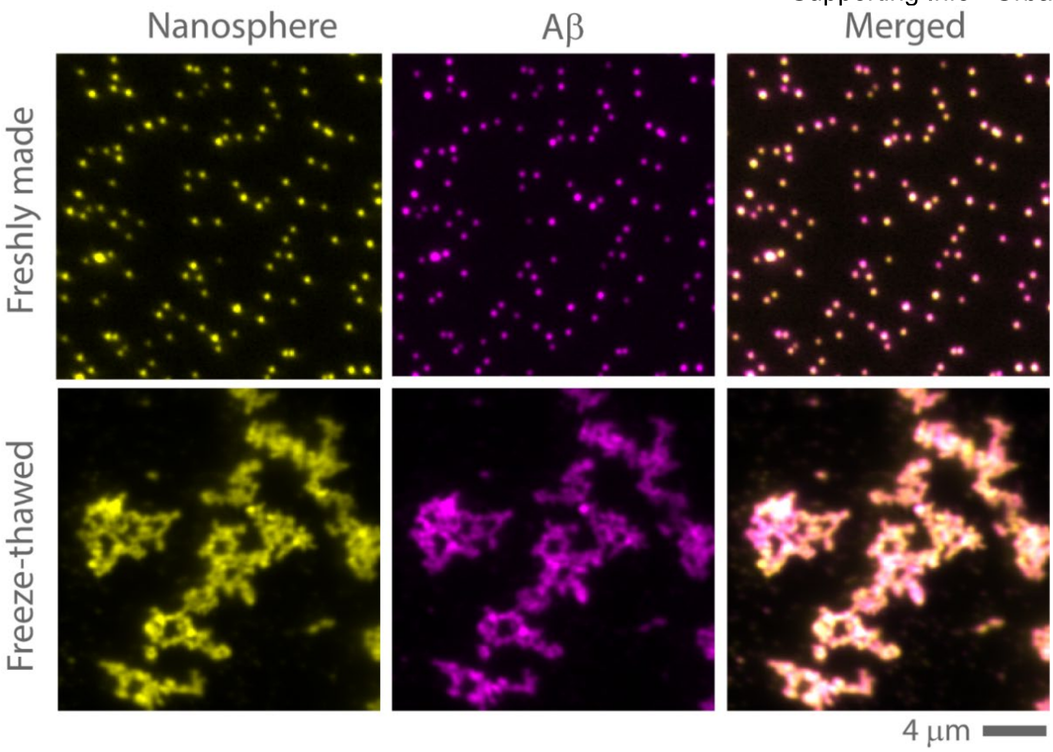


**Supplementary Figure 12.** SiMPull images illustrate Aβ42 aggregates on the surfaces of 30nm nanospheres from the same batch, both freshly prepared and after undergoing one freeze-thaw cycle. The freeze-thaw process leads to clumping of these nanospheres.

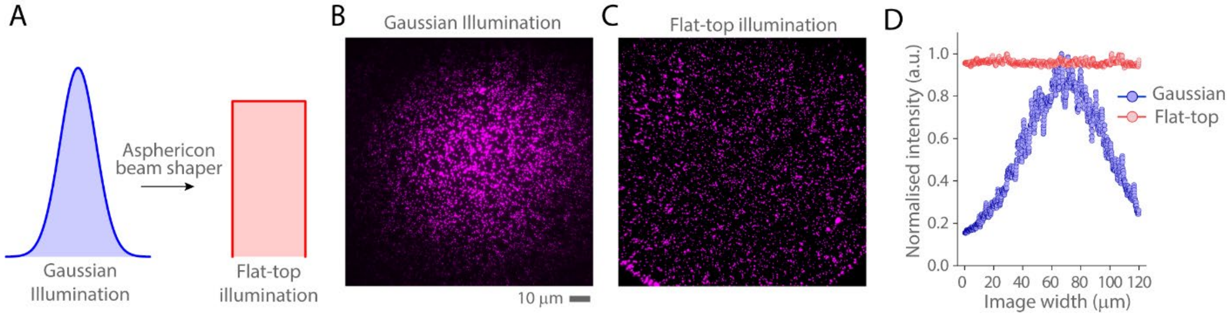


**Supplementary Figure 13. (A)** Conventional wide-field illumination with a gaussian profile is transformed to a flat-top profile by placing an Asphericon beam-shaping device in the excitation path. **(B-C)** show SiMPull images of Aβ42 aggregates engineered on the surfaces of 30nm nanospheres under **(B)** Gaussian illumination and **(C)** flat-top illumination, respectively. **(D)** The normalised integrated intensity profile of the 120x120 µm images is plotted for Gaussian (blue) and flat-top illumination (red). The intensity variation between the centre and edge of the image for Gaussian illumination exceeds 80%, whereas for flat-top illumination, this variation is less than 5%.

**Supplementary Table 1: Details of the post-mortem tissue used**

| Disease | Brain area | Age | Gender |
|---|---|---|---|
| AD | Frontal cortex | 92 | F |
| AD | Frontal cortex | 86 | M |
| AD | Frontal cortex | 79 | F |
| PD | Midbrain | 81 | F |
| PD | Midbrain | 75 | M |
| PD | Midbrain | 77 | F |

**Supplementary Table 2: Details of the Aβ aggregates used in this study**

| Protein | Vendor (Manufacturer) | Cat. No. | Measurements of aggregates | |
|---|---|---|---|---|
| | | | Conc. (μm) | Time (minutes) |
| **WT Aβ42** | Stratech (rPeptide) | A-1167-2 | 3 | 25 |
| **WT Aβ40** | Stratech (rPeptide) | A-1157-2 | 30 | 400 |
| **Aβ42 HiLyte™ Fluor 488-labeled + Aβ42 HiLyte™ Fluor 647-labeled** | Eurogentec (AnnSpec) | AS-64161<br>AS-60479-01 | 1.5 + 1.5 | 25 (for oligomer)<br>180 (for fibrils) |
| **Aβ40: Aβ42 9:1** | - | - | 27+3 | 120 |
| **Aβ40: Aβ42 7:3** | - | - | 21+9 | 260 |
| **E22G Aβ42 (Arctic)** | Eurogentec (AnnSpec) | AS-61967-05 | 3 | 20 |
| **E22G + WT Aβ42 1:1** | - | - | 1.5 + 1.5 | 10 |
| **E22Q Aβ42 (Dutch)** | Eurogentec (AnnSpec) | AS-62142 | 3 | 15 |
| **E22Q + WT Aβ42 1:1** | - | - | 1.5 + 1.5 | 5 |
| **Scrambled Aβ42 FAM labelled** | Eurogentec (AnnSpec) | AS-60892 | - | - |

**Supplementary Table 3: Details of the antibodies used in SiMPull study**

| Figure# | **Capture antibody** (Biotinylated) | Channel 1 (nm) Ex: 488 nm Em : 500-540 nm | Channel 2 Ex: 561 nm Em: 580-620 nm | Channel 3 Ex: 637 nm Em: 680-720 nm |
|---|---|---|---|---|
| **1 G-H, K-L** | 6E10 | Fluorescent nanosphere | Alexa 561 Fluor labelled A11 | Alexa 647 Fluor labelled 6E10 |
| **1 I-J, M-N** | 6E10 | Amytracker-540 | - | Alexa 647 Fluor labelled 6E10 |
| **4 F-G** | 6E10 | Fluorescent nanosphere | Alexa 561 Fluor labelled Aβ40 specific EPR23712-2 | Alexa 647 Fluor labelled Aβ42 specific 21F12 |

| | | | | |
|---|---|---|---|---|
| **5 F-G** | 6E10 | Fluorescent nanosphere | Alexa 561 Fluor labelled 4G8 | Alexa 647 Fluor labelled 6E10 |
| **6E-F** | MJFR1 | Fluorescent nanosphere | Alexa-561-fluor conjugated aSyn confirmation specific MJFR-14-6-4-2 antibody | Alexa 647 Fluor labelled MJFR1 antibody |
| **SI Fig3** | - | Fluorescent nanosphere | Alexa 561 Fluor labelled A11 | Alexa 647 Fluor labelled 6E10 |
| **SI Fig 10** | 6E10 | Fluorescent nanosphere | Alexa 561 Fluor labelled A11 | Alexa 647 Fluor labelled 6E10 |
| **SI Fig11, SI Fig 12** | 6E10 | Fluorescent nanosphere | - | Alexa 647 Fluor labelled 6E10 |
